# In silico design and computational characterization of novel chimeric multiepitope antigens for Mpox serosurveillance

**DOI:** 10.1101/2025.07.09.663824

**Authors:** Roland Ngwese Akwelle, Robert Adamu Shey, Mary Teke Efeti, Lekeayi Nelson Acha, Gordon Takop Nchanji, Tangan Yanick Aqua Stong, Joan Amban Chick, Ntang Emmaculate Yaah, Cabirou Mounchili Shintouo, Bernis Neneyoh Yengo, Abey Blessings Ayuk, Junior Ekunidi Engarimbi, Ketura Yaje Gwei, Brinate Teke Tebo, Derrick Neba Nebangwa, Muyer Tamnjong Beltine, Luc Vanhamme, Jacob Souopgui, Vincent P K Titanji, Stephen Mbigha Ghogomu

## Abstract

**Background:** The Monkeypox (Mpox) virus is a zoonotic Orthopoxvirus with a global outbreak that began in 2022 and spread to more than 128 countries, with more than 132,000 confirmed cases and 1500 deaths. The pandemic preparedness pipeline emphasizes the importance of diagnostic surveillance of pathogens in at-risk populations to monitor transmission and mitigate the impact on public health. Unfortunately, the current gold standard diagnostic tool for Mpox is limited in its field applicability. Therefore, there is a crucial need for the development of robust novel diagnostic tools to enable continuous surveillance of the disease. As such, this work sought to design and validate novel multiepitope antigens as diagnostic tools for Mpox serosurveillance.

**Methods:** Using immunoinformatic approaches, two novel Mpox multiepitope antigens (MP-MEDA-1 and MP-MEDA-2) were designed using linear B-epitopes of viral proteins previously characterized in Mpox serodiagnosis. The 3D structures of the designed antigens were predicted, refined, and validated. Protein-protein docking and interaction analyses were performed between the designed diagnostic antigens and the Fab regions of human IgA, IgG, and IgM.

**Results:** The designed antigens were predicted to be antigenic and demonstrated thermostability with desirable physicochemical properties. In addition, both antigens also demonstrated stable interactions with the Fab regions of selected immunoglobulins, with several residues interacting at the interfaces of all the docked complexes.

**Conclusions:** These preliminary findings highlight the potential of the MP-MEDA-1 and MP-MEDA-2 antigens as candidates to be further characterized for Mpox serosurveillance. The next phase of this project will focus on the expression and serological characterization of both antigens to determine their diagnostic parameters (sensitivity, specificity, and others).

## 1 Introduction

Mpox (previously known as monkeypox) is a zoonotic disease caused by the Mpox virus (MPXV), with sporadic outbreaks reported primarily in West and Central Africa (1). The predominant symptom of MPXV infection is skin rashes (often accompanied by anogenital or oropharyngeal/perioral mucosal lesions) (2). However, fever, chills, headache, and lymphadenopathy are also commonly reported symptoms in patients (3). The MPXV consists of 2 genetically distinct clades (clade I and clade II), with clade I (formerly called the Central African clade) infections associated with more severe disease and higher mortality (4, 5). On the other hand, clade II (formerly called the West African clade) is known to cause less severe disease with lower case fatality rates (CFRs).

From 2022, 2 major Mpox outbreaks have occurred in succession. A subclade of clade II (clade IIb), first identified in Nigeria in the 1970s, recently spread rapidly, causing a global outbreak in May 2022 (6). As a result, the World Health Organization declared the MPXV outbreak on 23 July 2022 a public health emergency of international concern (PHEIC) (7). This outbreak is estimated to have resulted in more than 100,000 cases and over 220 deaths in 122 countries, many of which had never previously reported the disease (8). In October 2023, a large-scale mpox outbreak caused by clade I occurred in the DR Congo, and in March 2024, the first individuals with mpox were reported outside endemic areas in the Democratic Republic of the Congo (9). As of 5 January 2025, more than 9,500 laboratory-confirmed cases were reported in the DR Congo, with an estimated CFR of 3.4% (10). However, the full scope and transmission characteristics of both outbreaks are unknown due to likely underestimation of the number of people infected, limited genomic surveillance, and human movements throughout central Africa (11).

The recent pattern of recurring outbreaks points to the essential need for increased medical and genomic surveillance of the monkeypox virus (12). This is particularly important in Africa, where concurrent outbreaks have been reported, especially in countries bordering the DR Congo, where the 2023 outbreak emerged (10), in addition to several other outbreaks (13). Genomic surveillance, together with corresponding epidemiological and clinical data, remains vital for assessing the epidemic trends in real-time and can be harnessed for the development of suitable countermeasures (including diagnostics, therapeutics, and vaccines) and the formulation of policies for effective prevention and control (14, 15).

Although genomic surveillance can generate vital data that can be employed for Mpox prevention and control, the genomic surveillance of MPXV (just like what happened for SARS-CoV-2) currently faces several major challenges (16). Despite the enormous endeavour, in reality, only a very small proportion of cases (from a few selected countries) have been sequenced, and many countries in Africa have very low sequencing capacity (17). Therefore, new variants could be circulating, but remain undetected owing to limited genomic surveillance capacity in several countries. In addition, complex transmission dynamics urgently necessitate increased genomic surveillance, especially considering the current extent of global connectivity and travel networks. Finally, recombination, which changes the viral genetic structure, has been reported in both clades of MPXV (18), with the MPXV clade Ib genomes diverging at a faster rate than those of clade IIb (2022), primarily due to an unusually high incidence of recombination (19).

Furthermore, integrated surveillance incorporating patient testing, pathogen genomic sequencing, population serosurveillance, and mortality tracking has been reported to be essential for epidemic preparedness (20). Additionally, the occurrence of asymptomatic cases (estimated to occur in 1.8 to 6.5% of persons with Mpox infection) is reported to contribute to the underestimation of the disease burden and warrants the use of seroprevalence surveys (21). In addition to genomic surveillance for emerging infectious pathogens, serosurveillance has been a key approach for measuring population exposure and immunity to a pathogen and understanding the potential for pathogen escape (22). Serological techniques enable real-time population immunity assessment and may enable the identification of circulating strains based on their immunological signatures in longitudinal surveys (20). Serosurveillance studies, therefore, have great potential to enhance the understanding of the current Mpox outbreak, particularly in the African context.

Although the detection of viral DNA by PCR is the gold standard to confirm MPXV infection, the sophistication of this technique limits its applicability in field settings (23). Antibodies against MPXV can be detected in serum approximately 1–2 weeks after the onset of symptoms and can aid in diagnostics under certain circumstances (24). However, despite several attempts, there are no approved serological tests to quantify humoral immune responses to MPVX infection. Nevertheless, different studies have reported the role of IgA, IgG, and IgM as serological markers associated with the acute and post-acute phases of infection (25, 26, 27). With the persistent transmission and evolution of MPXV, disease symptoms have become milder, with some infections being asymptomatic (despite ongoing transmission) (28). In addition, since the 2022 and 2023 outbreaks, MPXV has become more contagious (with sustained and efficient human-to-human transmission) (29). Hence, there is an urgent need for continuous research for robust diagnostic and screening tools that can be deployed for Mpox serosurveillance, as these tools will be important to prevent and control MPXV epidemics (30). Immunoinformatics-based approaches have been deployed as a suitable strategy to design multiepitope chimeric antigens with desirable diagnostic parameters (31). Multi-epitope diagnostic antigens with increased sensitivity and specificity have been designed for the serodiagnosis of several diseases, including onchocerciasis (32, 33, 34), tuberculosis (35), and Chagas (36, 37). This work is focused on the computational design and characterization of two novel multi-epitope diagnostic antigens (MP-MEDA-1 and MP-MEDA-2) using immunoinformatic approaches.

## 2 Materials and methods

### 2.1 Protein sequence retrieval and preliminary characterization

The sequences of 5 MPVX (strain Zaire-96-I-16) proteins (B21R, N3R, A30L, E8L, and B18R with accession numbers Q8V4Q7, Q8V4Q2, Q8V4U9, Q8V4Y0, and Q8V4R0, respectively), which were previously assessed for their diagnostic potential were retrieved from the UniProtKB database in the *.fasta* format for preliminary analyses as previously described (38, 39). The initial characterization included principally signal peptide prediction using the SignalP 6.0 (https://services.healthtech.dtu.dk/services/SignalP-6.0/) and TOPCONS (https://topcons.cbr.su.se/) servers, as well as transmembrane domain prediction using the DeepTMHMM 1.0 (https://services.healthtech.dtu.dk/services/DeepTMHMM-1.0/) and TOPCONS (https://topcons.cbr.su.se/) servers. The SignalP 6.0 server is based on machine-learning protein language models (LMs) and uses information from millions of unannotated protein sequences across all domains of life to predict all five signal peptides (SP) types (40). The DeepTMHMM 1.0 server employs a deep learning protein language model-based algorithm to detect and predict the topology of both alpha-helical and beta-barrel proteins with extraordinary accuracy (41). The TOPCONS server is a consensus-based membrane protein topology and signal peptide prediction tool with state-of-the-art performance in topology predictions (42).

### 2.2 Linear B-epitope prediction

Linear B-lymphocyte epitope prediction was performed using the BepiPred-3.0 (https://services.healthtech.dtu.dk/service.php?BepiPred-3.0) and LBtope (https://webs.iiitd.edu.in/raghava/lbtope/index.php) web servers. The BepiPred-3.0 web server is a sequence-based tool that uses numerical representations from protein language model (LM) embeddings to significantly improve the accuracy of both linear and conformational B-cell epitope prediction on many independent test sets (43). Epitope prediction analysis on the BepiPred-3.0 server for the protein sequences was conducted using the server’s default threshold of 0.1512. The LBtope server, on the other hand, is an epitope prediction tool that uses a larger dataset of experimentally validated B-cell epitopes and non-B-cell epitopes to predict epitopes based on machine learning techniques. Using diverse features such as binary profiles, dipeptide compositions, and amino acid pair (AAP) profiles, the server can achieve accuracies from ∼54% to 86% (44).

### 2.3 Epitope antigenicity prediction

To design multiepitope chimeric diagnostic antigens for serological studies, the VaxiJen v2.0 server (http://www.ddg-pharmfac.net/vaxijen/VaxiJen/VaxiJen.html) was used to predict the antigenicity of all the predicted epitopes. The VaxiJen v2.0 server is the first alignment-independent approach that uses the physicochemical characteristics of peptides and proteins for antigen prediction. The server algorithm deploys the autocross-covariance (ACC) transformation of input peptide sequences into uniform vectors of principal amino acid features to predict the antigenicity of bacterial, fungal, parasitic, viral, and tumor peptides/proteins (45). The default threshold of 0.4 for the virus model was used to predict the antigenicity of each predicted epitope.

### 2.4 Multiepitope diagnostic antigen design

The selected antigenic linear-B epitopes were combined into two multiepitope diagnostic antigens (designated MP-MEDA-1 and MP-MEDA-2) via GSGSG flexible linkers. Each antigen had a 10×His tag added to its N-terminal region. The 10×His tags were followed by the HRV 3C protease cleavage sequence (LEVLFQ/GP) and the different peptide sequences separated by GSGSG linkers. The GSGSG linker has been reported to maximize epitope stability and recognition (46) and has previously been used in the design of multiepitope diagnostic antigens for other infectious diseases, including onchocerciasis (33, 34), Chagas disease (47), and hepatitis B (48).

### 2.5 Physicochemical Properties and Solubility Prediction

The physicochemical properties of the designed multiepitope diagnostic antigens were predicted using the ProtParam web server (https://web.expasy.org/protparam/). The ProtParam server generates information on different protein parameters, including molecular weight, aliphatic index, theoretical pI, instability index, amino acid, and atomic composition. In addition, the server also predicts the half-life of the input protein sequence upon expression in *Escherichia coli*, yeast, and mammalian cells (49). The solubilities of the designed antigens upon expression in bacteria were predicted using the NetSolP-1.0 (https://services.healthtech.dtu.dk/services/NetSolP-1.0/) and DeepSoluE (http://lab.malab.cn/∼wangchao/softs/DeepSoluE/) servers. The NetSolP-1.0 server deploys a working algorithm based on deep-learning protein language models to predict the solubility and usability of proteins expressed in *Escherichia coli* directly from the sequence. The server achieves state-of-the-art performance and outperforms current tools used for in-silico protein solubility and usability prediction (50). The DeepSoluE server, on the other hand, is a novel tool that predicts protein solubility using a long short-term memory (LSTM) network with hybrid features comprising physicochemical properties and distributed representations of amino acid residues (51).

### 2.6 Multiepitope diagnostic protein antigenicity prediction

The antigenicity scores of the designed multi-epitope chimeric antigens were predicted using the VaxiJen v2.0 (described above) and the ANTIGENpro server from the Scratch Proteomics Predictor platform (https://scratch.proteomics.ics.uci.edu/). The ANTIGENpro server, similar to the VaxiJen v2.0 server, is a sequence-based, alignment-free prediction tool trained on a large, non-redundant dataset obtained from protein microarrays, achieving an accuracy of 82% (52).

### 2.7 Secondary structure and intrinsic disorder prediction

The PSIPRED v4.0 (http://bioinf.cs.ucl.ac.uk/psipred) and NetSurfP-3.0 (https://services.healthtech.dtu.dk/services/NetSurfP-3.0/) servers were used to predict the secondary structures of the designed chimeric antigens. PSIPRED is a simple and accurate protein secondary structure prediction approach that incorporates two feed-forward neural networks that perform an analysis on the outputs obtained from a position-specific iterated BLAST (PSI-BLAST) (53). The current version of the PSIPRED server (v4.0) has a Q3 prediction accuracy of 84.2.5% (54). Moreover, the NetSurfP–3.0 server uses convolutional and long short-term memory neural networks to generate cutting-edge predictions for protein secondary structure, solvent accessibility, disorder, and backbone geometry from protein sequences while exploiting recent advances in pretrained protein language models to drastically improve runtime (55). The server’s accuracy has been tested on several independent test datasets and found to consistently produce accurate predictions for each of its output features (56).

Since unfolded proteins have been identified as an important class of structural antigens (57) that can be recognized by antibodies (58), intrinsically disordered regions (IDRs) in the designed chimeric diagnostic antigens were predicted using the AIUPred and flDPnn2 web servers. The AIUPred server (https://aiupred.elte.hu/) is a novel version of the previously reported IUPred server (59) that incorporates deep learning techniques into the energy estimation framework, achieving improved performance (60). On the other hand, the flDPnn2 server (https://biomine.cs.vcu.edu/servers/flDPnn2/) uses deep neural networks to provide accurate and fast predictions of intrinsic disorder using protein sequences (61).

### 2.8 Solvent accessibility and thermal stability prediction

The relative solvent accessibilities for both designed antigens were predicted using the E-pRSA (https://e-prsa.biocomp.unibo.it/main/) and the DeepREx-WS web (https://deeprex.biocomp.unibo.it/home/) servers. The E-pRSA server is a novel approach to estimating Relative Solvent Accessibility values (RSAs) of residues directly from a protein sequence. The method exploits two complementary Protein Language Models (ProtT5 and ESM2) to provide rapid and precise predictions (62). The DeepREx-WS server, on the other hand, is a tool that provides a multidimensional characterization of exposed and buried residues in a protein using sequence information. The server uses a novel deep learning-based method, DeepREx, for the two-class prediction of protein solvent exposure and has been trained and tested on high-quality structures extracted from the PDB (63).

Since the cold chain is usually a key requirement for most biologics needed for infectious disease control, thermal stability predictions for both designed antigens were performed using the DeepSTABp server (https://csb-deepstabp.bio.rptu.de/) and the TemStaPro tool. The TemStapPro tool was accessed through the NeuroSnap platform (https://neurosnap.ai/service/TemStaPro%20Protein%20Thermostability%20Prediction). DeepSTABp performs end-to-end protein melting temperature prediction by deploying a transformer-based protein language model for sequence embedding and state-of-the-art feature extraction in combination with other deep learning techniques (64). TemStaPro, on the other hand, functions on the basis of a large-scale, comprehensive approach to using protein language model (pLM) embeddings to predict protein stability above certain temperature thresholds (65).

### 2.9 Tertiary structure prediction, refinement, and validation

The ColabFold platform, which offers an easy-to-use interface to use the AlphaFold2 (66) technology within the Google Colab environment, was used to predict the 3D structure of the designed diagnostic antigens. ColabFold combines the quick homology search of MMseqs2 with AlphaFold2 or RoseTTAFold to provide faster protein structure and complex prediction (67). ColabFold is an innovative protein prediction tool designed to facilitate structure modeling for users. As a result, it provides pre- and postprocessing steps as well as an intuitive interface for several protein prediction models (68). The 3D structures obtained were then refined using a two-step procedure, in the first step using the freely accessible FG-MD web server (https://seq2fun.dcmb.med.umich.edu/FG-MD/). The FG-MD server deploys a molecular dynamics (MD)-based algorithm for atomic-level protein structure refinement. The FG-MD refinement algorithm combines the physical-based force field AMBER99 with knowledge-based hydrogen bonding and repulsive potentials and demonstrates significant potential in atomic-level protein structure refinements when tested on both benchmark and CASP refinement experiments (69). A further refinement step was performed on the GalaxyRefine server (http://galaxy.seoklab.org/cgi-bin/submit.cgi?type=REFINE). The GalaxyRefine server is based on a method that first rebuilds side chains and then performs side-chain repacking and subsequent overall structure relaxation using molecular dynamics simulation. The method was evaluated in CASP10 community-wide tests and showed the best performance in improving the local structure quality (70). The ProSA-web (https://prosa.services.came.sbg.ac.at/prosa.php) and ERRAT (http://services.mbi.ucla.edu/ERRAT/) web servers were used to assess the overall quality of the modeled 3D structures of MP-MEDA-1 and MP-MEDA-2. The ProSA tool is widely deployed to inspect 3D models of protein structures for potential errors and is used to detect errors in experimentally determined structures and theoretical models (71). The ERRAT server, on the other hand, analyzes non-bonded atomic interactions in the modeled 3D structure in comparison to more reliable high-resolution structures obtained by crystallography (72). Finally, a Ramachandran plot analysis to determine the quality of the refined 3D structures based on the percentage of residues in allowed and disallowed regions was performed using the PDBsum server (https://www.ebi.ac.uk/thornton-srv/databases/pdbsum/Generate.html).

### 2.10 Discontinuous B-cell epitope prediction

After validating the 3D structures of the designed diagnostic antigens, the ElliPro server (http://tools.iedb.org/ellipro/) was used to predict discontinuous B-cell epitopes using the default parameters (minimum score: 0.5; maximum distance: 6 Å). The server implements a previously developed method that represents protein structures as ellipsoids and performs calculations of protrusion indices for residues outside the ellipsoid. The ElliPro tool was tested on a benchmark dataset of discontinuous epitopes inferred from 3D structures of antibody-protein complexes. Compared to six other structure-based epitope prediction methods, ElliPro achieved the best performance, with an AUC value of 0.732 when the most significant prediction for each protein was considered (73).

### 2.11 Protein‒protein docking, binding affinity, and interaction analyses

The structures of the Fab regions of human IgA, IgG, and IgM (PDB IDs: 7K75, 7K76, and 1DEE, respectively) were downloaded from the Research Collaboratory for Structural Bioinformatics (RCSB) Protein Data Bank (PDB) database (https://www.rcsb.org/) for protein-protein interaction analyses. The RCSB PDB is a protein structure database that facilitates access to annotated data on the three-dimensional (3D) structures of macromolecules (including proteins and nucleic acids) and related small compounds (including drugs, cofactors, and inhibitors) in the PDB repository, thereby supporting scientific research and teaching globally (74). The RCSB PDB website provides tools to search, visualize, and analyze PDB data (75). IgA, IgG, and IgM antibodies have been implicated in disease onset and progression and have been investigated in serological assay development (25, 26, 76). The protein structures were then cleaned by removing water molecules and unnecessary structures using the PyMOL tool. PyMOL is a Python-based cross-platform molecular graphics software widely used for three-dimensional (3D) visualization of proteins, nucleic acids, small molecules, electron densities, surfaces, and trajectories. The tool is also efficient for editing molecules, ray tracing, and making movies (77). To assess interactions between the designed multiepitope diagnostic antigens and the selected immunoglobulins, protein‒protein docking analyses were performed on the ClusPro 2.0 web server (https://cluspro.bu.edu/login.php). The “Antibody mode” was selected during docking, with the immunoglobulin Fab regions selected as receptors and the antigens selected as ligands. The ClusPro 2.0 server has consistently been ranked among the top docking servers, offering excellent predictive performance for protein‒protein complex docking according to the Critical Assessment of Predicted Interactions (CAPRI) (78). The following computational procedures are used by the server to accomplish docking: (i) rigid body docking by sampling billions of conformations; (ii) clustering of the 1000 lowest-energy structures produced using root mean square deviation (RMSD) to identify the largest clusters that represent the most likely models of the complex; and (iii) energy minimization for refinement of selected structures. PIPER, a docking program based on the Fast Fourier Transform (FFT) correlation technique, is used in the rigid body docking step (79). To perform similarity-based docking, the server retrieves templates from a database of experimentally determined structures and builds models using energy-based optimization, permitting structural flexibility (78).

The binding affinities for each of the predicted complexes were analyzed using the Protein Binding Energy (PRODIGY) prediction server (https://wenmr.science.uu.nl/prodigy/) as previously described (80). The PRODIGY server implements a simple but very robust predictive model based on intermolecular interactions and features derived from the non-interface surface (81). In addition, the AREA-AFFINITY web server, which performs binding affinity prediction for protein‒protein or antibody‒protein interactions based on interface and surface areas in the structure of protein‒protein complexes, was also used (82). The docked complexes were then further analyzed on the PDBsum server (http://www.ebi.ac.uk/thornton-srv/databases/pdbsum/Generate.html) and the PICKLUSTER tool on ChimeraX (v1.9). The PDBsum server generates 2D pictures of the interactions at the ligand‒protein interface from the submitted 3D coordinates (83). The PICKLUSTER [Protein Interface C(K)luster] is a UCSF ChimeraX plug-in that clusters protein interfaces based on distance (84). Interface calculation was done with clustering, with the default cutoff for interface clustering set to 5 and the pLDDT cutoff for interface calculation set to 50.

### 2.12 Molecular dynamics (MD) simulations

Identifying flexible areas of protein structures is essential for understanding their biological roles (85). Normal-mode analysis (NMA) for the docked diagnostic antigen-Ig Fab complexes was performed via the iMODS server (http://imods.Chaconlab.org/). The server provides several representations of effective motion, including a vector field, affine-model arrows, and modal animations. In addition, a detailed analysis includes profiles of mobility (NMA B-factors), covariance maps, deformability, eigenvalues, and linking matrix (86). The iMODS server has generated results that are consistent with targeted molecular dynamics and crystallographic anisotropic displacement studies (86). The analyses of the docked antigen-Ig Fab complexes were executed to ascertain the internal dihedral coordinates while simultaneously assessing the cooperative functional movements. Specific parameters, including B-factors, complex deformability, eigenvalues, covariance values, and elastic models of the docked complexes, were predicted using the server. In addition, the AGGRESCAN3D v. 2.0 server (http://biocomp.chem.uw.edu.pl/A3D2/) was used to predict the solubility and aggression propensity of each of the complexes. The server integrates the 3D information of protein structures and evaluates the contribution of solvent-exposed aggregation-prone regions. The algorithm functions by projecting experimental aggregation propensities onto a protein structure (87).

### 2.13 Codon optimization and in-silico cloning

Since bacterial expression is often preferred as the first line for protein expression experiments (88), the K12 *E. coli* strain was selected as a suitable host to express the multiepitope diagnostic antigens. Reverse translation of the protein sequences into DNA and codon optimization were achieved using the Java Codon Adaptation Tool (https://www.jcat.de/). The JCat server offers the flexibility to adapt codon usage for any user-provided sequence to that of any selected host organism to increase heterologous protein production (89). Calculations on the server are performed using an iterative algorithm that works based on a precise mathematical formulation (90). The server also generates data on the GC% content and codon adaptation index (CAI), both of which have been reported to influence protein expression (91). To express the designed antigens in bacteria, in-silico cloning was performed, and the codon-optimized gene sequences were inserted between the NdeI and XhoI restriction sites in the pET30a+ vector using the Snapgene software (https://www.snapgene.com/).

### 2.14 RNA secondary structure and tertiary structure prediction

The secondary and tertiary structures of RNA have been reported to affect stability and function (92), and can also impact protein expression as well as protein‒RNA interactions (93, 94). The codon-optimized DNA sequences coding for MP-MEDA-1 and MP-MEDA-2 were transcribed into mRNA using the Transcription and Translation Tool (https://biomodel.uah.es/en/lab/cybertory/analysis/trans.htm). The RNAfold web server (http://rna.tbi.univie.ac.at/cgi-bin/RNAWebSuite/RNAfold.cgi) was used (with the default parameters) to predict the secondary structures of the mRNA sequences. The RNAfold server performs the minimum free energy (MFE) structure prediction of a single sequence using the classic algorithm of Zuker and Stiegler and can also calculate equilibrium base pairing probabilities via John McCaskill’s partition function algorithm (95). The trRosettaRNA server (https://yanglab.qd.sdu.edu.cn/trRosettaRNA/) was used to predict the 3D structure of the mRNA sequences coding for the designed antigens. The server algorithm is based on an automated deep learning-based approach to RNA 3D structure prediction. The trRosettaRNA pipeline begins with 1D and 2D geometries prediction using a transformer network in the first step, and then, tertiary structure folding by energy minimization (96).

## 3 Results

### 3.1 Protein Retrieval and Preliminary Characterization

The sequences of 5 Mpox virus proteins previously reported to be implicated in Mpox diagnosis, B21R (97), N3R (98), A30L (99), E8L (100), and B18R (100) (having accession numbers Q8V4Q7, Q8V4Q2, Q8V4U9, Q8V4Y0, and Q8V4R0, respectively) were downloaded from the UniProtKB database for preliminary analyses and epitope prediction. Signal peptide analyses predicted the presence of signal peptide sequences only in the B21R and N3R antigens. The signal peptide prediction results from the SignalP 6.0 server correlated with those from the TOPCONS server (Table 1). For TM domain prediction analyses, the TOPCONS server predicted the presence of TM regions in 3 of the 5 antigens, whereas 2 antigens were predicted to contain TM regions on the DeepTMHMM 1.0 server. For the combined prediction, the number of TM domains ranged from 1 - 3. Both servers predicted the presence of TM domains in Q8V4Q7 and Q8V4Y0, whereas Q8V4U9 was predicted to contain TM domains only by the TOPCONS server. However, the servers showed disparities in the number of TM domains for Q8V4Q7, with the DeepTMHMM 1.0 server predicting 3 TM domains in contrast to the 1 TM domain predicted by the TOPCONS server (Table 1).

**Table 1:** Preliminary characterization of antigens selected for epitope prediction.

| Protein | TM domain prediction |  | Signal peptide prediction |  |
| --- | --- | --- | --- | --- |
|  | TOPCONS | DeepTMHMM 1.0 | SignalP 6.0 | TOPCONS |
| Q8V4Q7 | 1 | 3 | Present | Present |
| Q8V4Q2 | 0 | 0 | Present | Present |
| Q8V4U9 | 1 | 0 | Absent | Absent |
| Q8V4Y0 | 1 | 1 | Absent | Absent |
| Q8V4R0 | 0 | 0 | Absent | Absent |

### 3.2 Linear B-cell epitope and epitope antigenicity prediction

The LBtope and BepiPred 3.0 servers were used to predict linear B-lymphocyte epitopes (with different numbers of amino acid residues), and epitopes concurrently predicted by both servers were selected to design the multiepitope chimeric diagnostic antigens. Following epitope prediction, the antigenicity of each epitope was predicted using the VaxiJen v2.0 server. A total of 23 linear B-epitopes with high antigenicity scores (the default threshold is 0.4) were selected to design two chimeric antigens (MP-MEDA-1 and MP-MEDA-2) (Table 2).

**Table 2:**
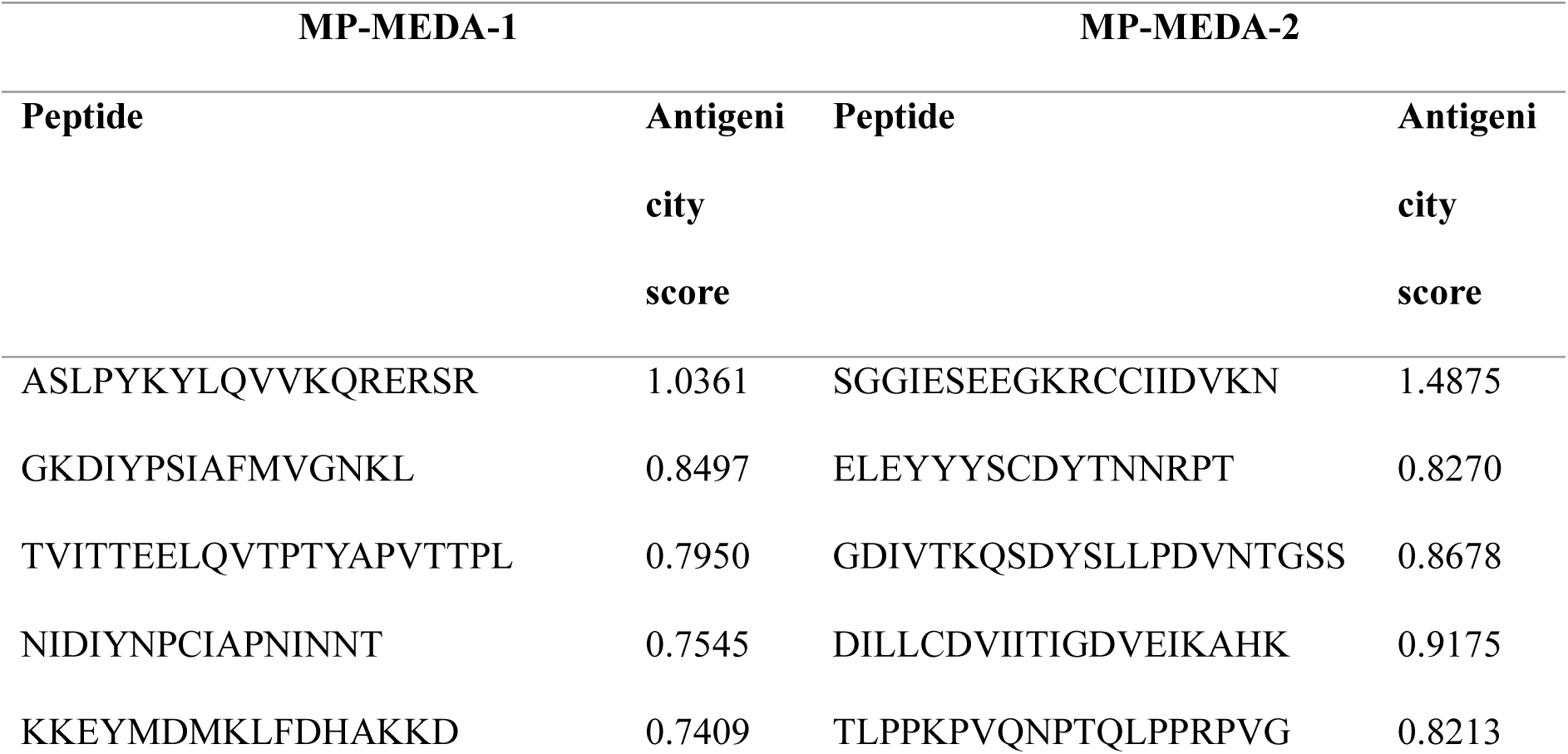
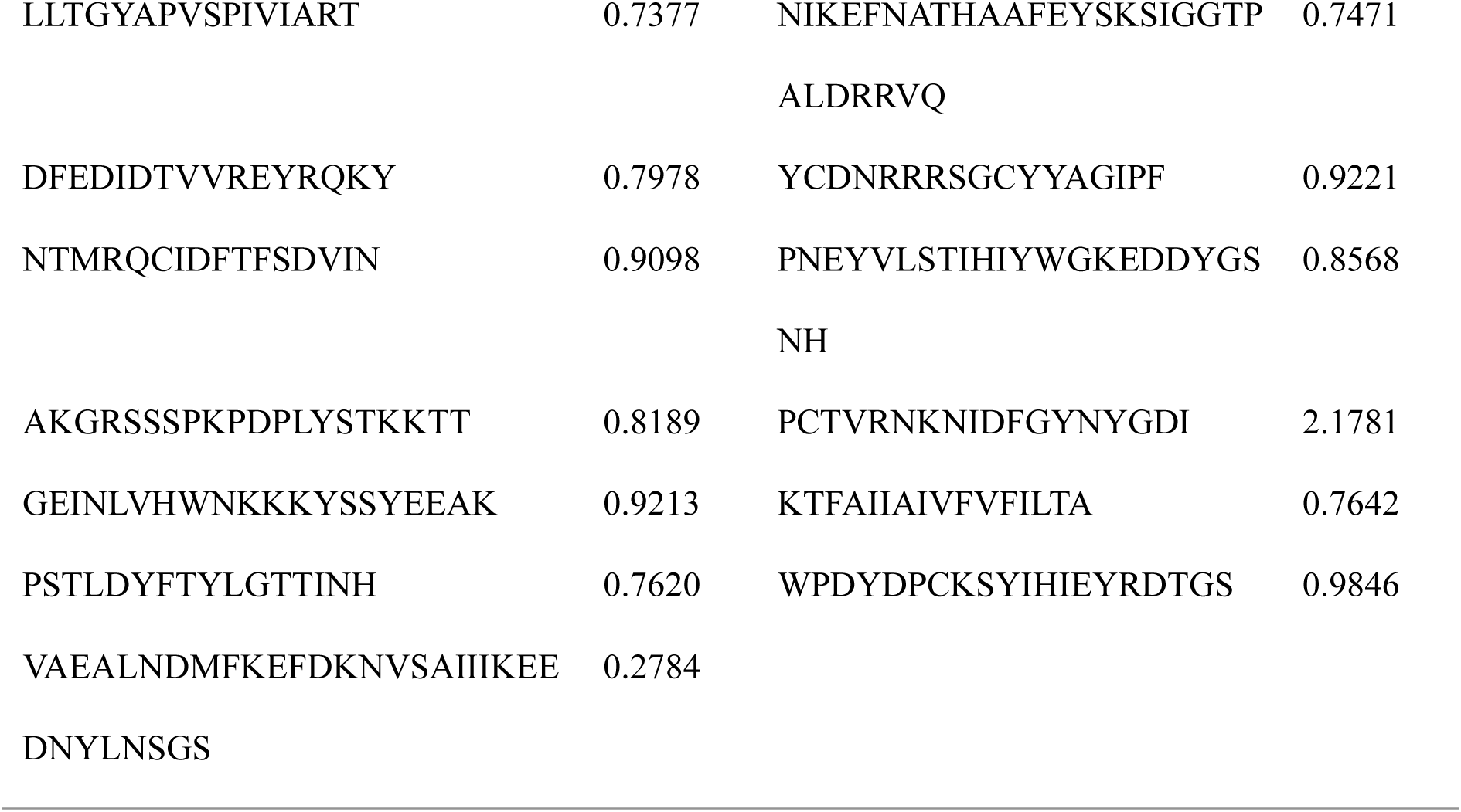
Predicted antigenicity scores of linear B-cell epitopes in MP-MEDA-1 and MP-MEDA-2.

### 3.3 Design of the Chimeric Multi-Epitope Diagnostic Antigens

The predicted epitopes with high antigenicity scores (above the default threshold of 0.4) were combined via GSGSG flexible linkers to design two multiepitope chimeric antigens (MP-MEDA-1 and MP-MEDA-2), with MP-MEDA-1 comprising 12 linear B-cell epitopes and MP-MEDA-2 comprising 11 linear B-cell epitopes. In addition, 10×His tags were added at the N-terminal regions of both chimeric antigens (to facilitate purification and identification) and separated from the epitopes using the HRV 3C protease sequence (LEVLFQ/GP). In summary, MP-MEDA-1 comprises 308 amino acid residues with a molecular weight of 32.6 kDa, whereas MP-MEDA-2 comprises 294 amino acid residues with a molecular weight of 30.9 kDa (Fig. 1).

**Figure 1.**
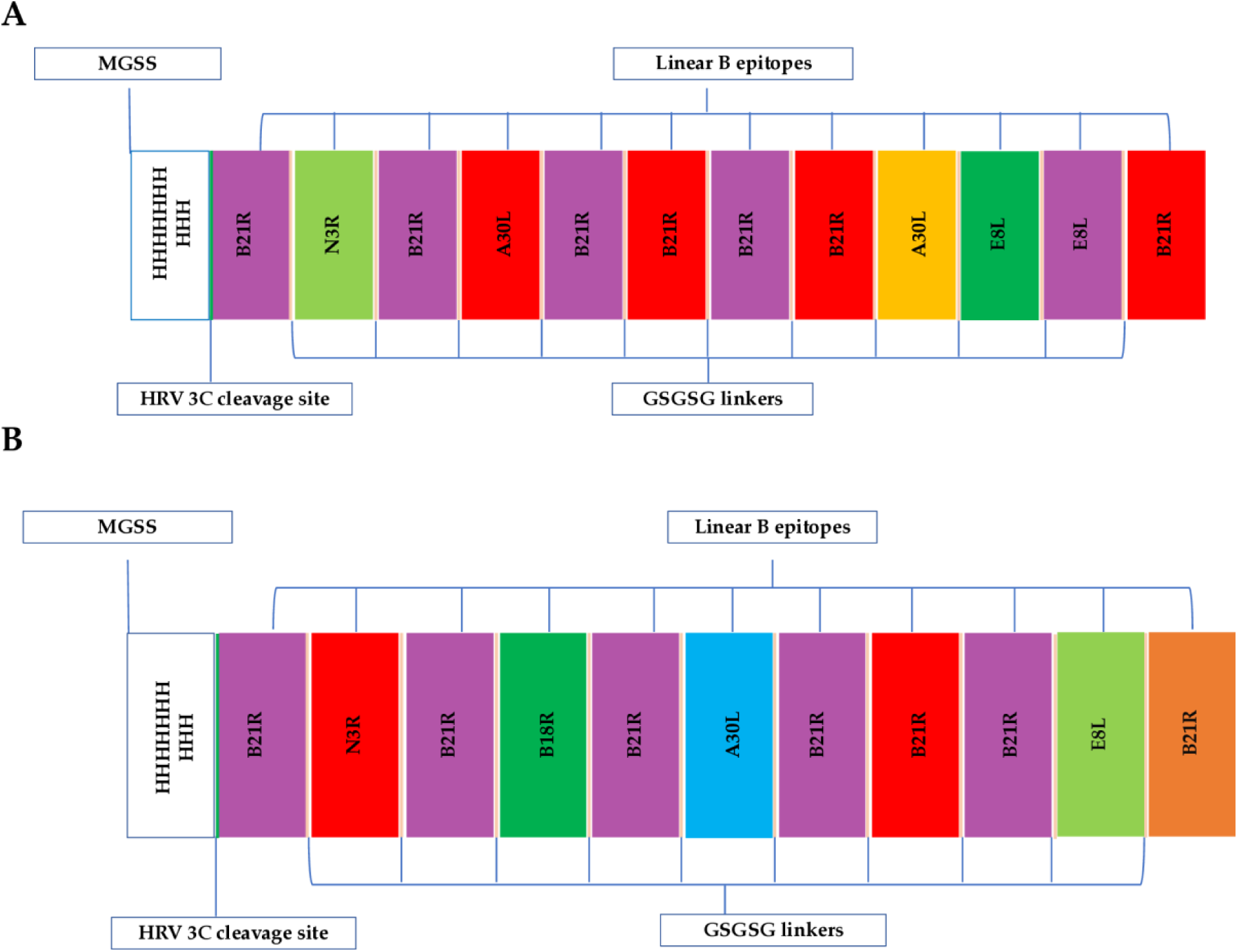
Schematic representation of the designed multiepitope diagnostic antigens, MP-MEDA-1 and MP-MEDA-2. **A.** The 308-amino-acid-long MP-MEDA-1 sequence, containing 12 linear B-cell epitopes connected using the GSGSG linker (cyan). **B**. The 294-amino-acid-long MP-MEDA-2 sequence, containing 11 linear B-cell epitopes connected using the GSGSG linker (cyan). Each antigen has an N-terminal 10×His tag (for purification and identification), followed by the HRV 3C cleavage sequence (green), and the different epitopes.

### 3.4 Physicochemical Properties and Solubility Analysis

The MP-MEDA-1 chimeric antigen, composed of 308 amino acids, was predicted to have a molecular weight (MW) of 32.6 kDa and a theoretical pI of 7.25, indicating that the protein is slightly basic. On the other hand, the MP-MEDA-2 antigen, comprising 294 amino acids, was predicted to have a molecular weight of 30.9 kDa and a theoretical pI of 6.08, indicating that the protein is moderately acidic. Both proteins displayed the same predicted half-life of 30 h in vitro in mammalian reticulocytes, >20 h in yeast, and >10 h in vivo in *E. coli*. The predicted instability index (II) was 30.40 for MP-MEDA-1 and 42.32 for MP-MEDA-2. These scores classify MP-MEDA-1 as stable and MP-MEDA-2 as slightly unstable (the threshold score is 40.00). The predicted grand average of hydropathicity (GRAVY) was -0.585 for MP-MEDA-1 and -0.525 for MP-MEDA-2, indicating that both proteins are hydrophilic and can interact with water molecules. The predicted aliphatic indices were 59.81 for MP-MEDA-1 and 60.99 for MP-MEDA-2, suggesting that both proteins have the potential to exhibit thermostability. The suggested thermal stability results are in line with the predicted melting temperature (*T*_m_) values of 46.6 °C for MP-MEDA-1 and 42.9 °C for MP-MEDA-2 on the DeepSTABp server. The results from the TemStaPro tool revealed a reduction in thermal stability for both proteins, with the average temperature increasing from 40 °C to 65 °C, with MP-MEDA-2 displaying a better stability profile across the different temperatures (Fig. S1, Table S1). The predicted solubility scores were 0.6556 for MP-MEDA-2 and 0.5481 for MP-MEDA-2 on the NetSolP 1.0 server. In addition, the predicted solubility scores on the DeepSoluE server were 0.7254 for MP-MEDA-2 and 0.7953 for MP-MEDA-2. The solubility scores predicted by the two servers suggest that both designed chimeric antigens are soluble upon expression in bacteria (Table 3).

**Table 3:**
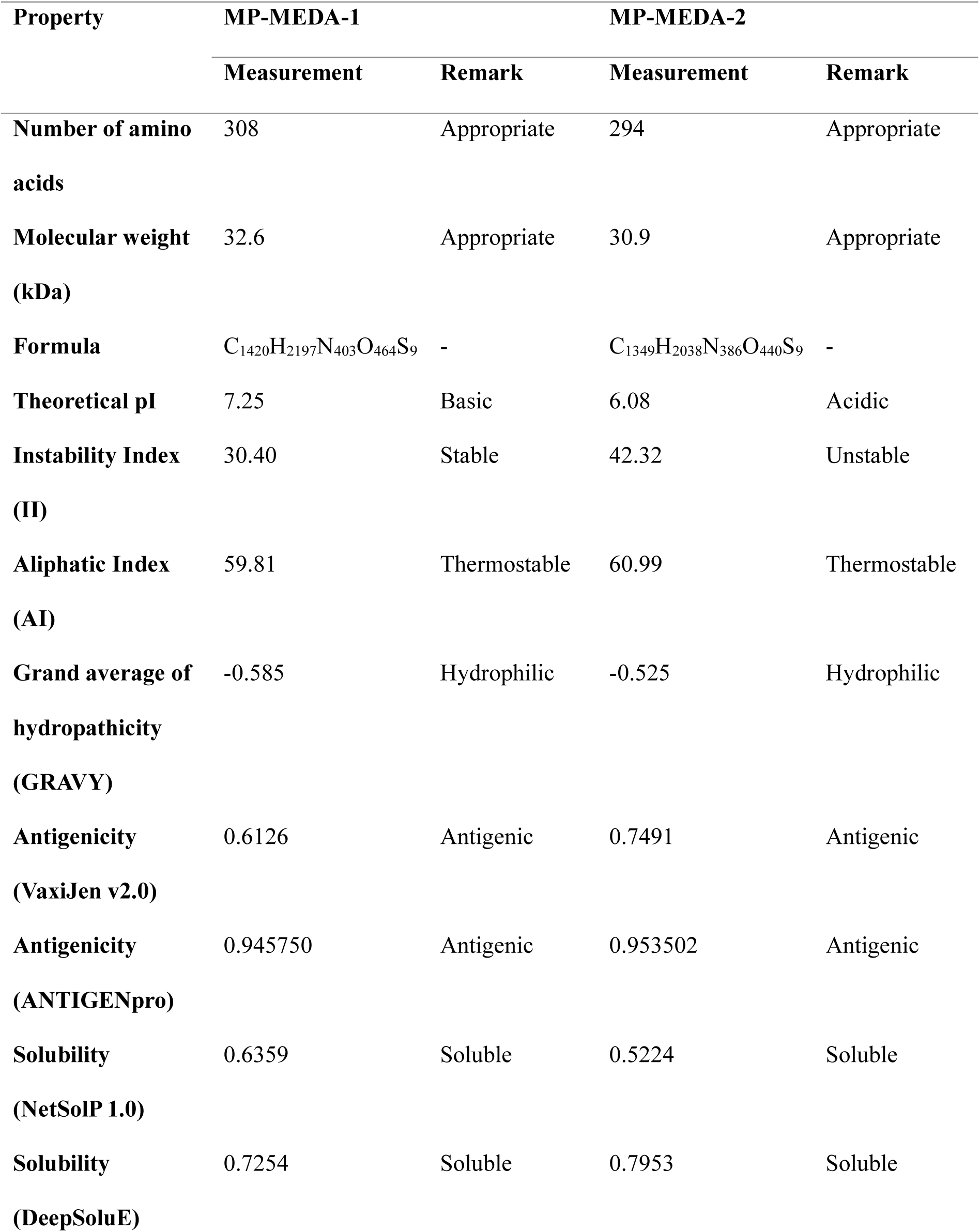
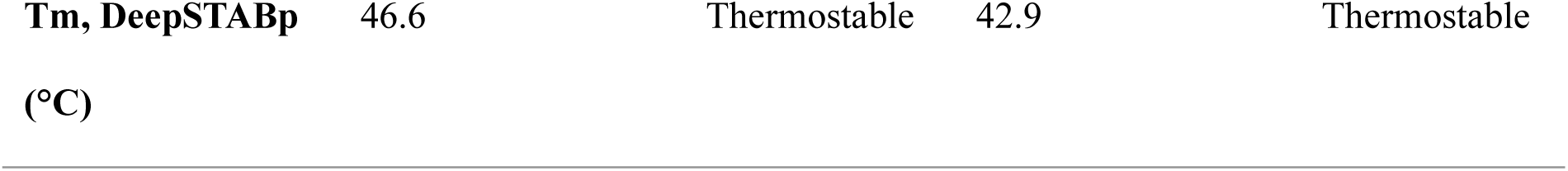
Physicochemical properties of the designed multiepitope diagnostic antigens (MP-MEDA-1 and MP-MEDA-2)

### 3.5 Antigenicity and thermal stability prediction

The VaxiJen v2.0 and ANTIGENpro servers were used to predict antigenicity scores for both multiepitope diagnostic antigens. The predicted antigenicity scores were 0.6126 for MP-MEDA-1 and 0.7491 for MP-MEDA-2 on the VaxiJen v2.0 server. In addition, the predicted antigenicity scores were 0.945750 for MP-MEDA-1 and 0.953502 for MP-MEDA-2 on the ANTIGENpro server. The predicted scores by both servers suggest that the designed chimeric antigens are antigenic, with MP-MEDA-2 having a slightly higher antigenic score than MP-MEDA-1 (Table 3).

### 3.6 Secondary structure, solvent accessibility, and intrinsic disorder prediction

The secondary structures of the multi-epitope antigens were predicted using the PSIPRED and NetSurfP-3.0. MP-MEDA-1 was predicted to consist of 10.0% alpha helices, 19.2% beta strands, and 70.8% coils. Moereover, MP-MEDA-2 was predicted to consist of 11.2% alpha helices, 22.5% beta strands, and 66.3% coils (Fig. 2). Considering solvent accessibility, MP-MEDA-1 was predicted to contain 303 (98.38%) amino acid residues exposed (RSA>=20%), whereas 5 (1.62%) residues were predicted to be buried (RSA<20%). In addition, 249 (80.84%) were predicted to be found at interaction sites. For MP-MEDA-2, 284 (96.6%) amino acid residues were predicted to be exposed (RSA>=20%), while 10 (3.4%) residues were predicted to be buried (RSA<20%). Compared with MP-MEDA-1, which has 249 residues (80.84%), MP-MEDA-2 was predicted to have fewer residues at interaction sites, with 144 (48.98%) residues predicted to be found at interaction sites (Fig. S2A, B, C, and D). Both proteins were also predicted to contain intrinsically disordered regions (Fig. S2E, F, G, and H).

**Figure 2.**
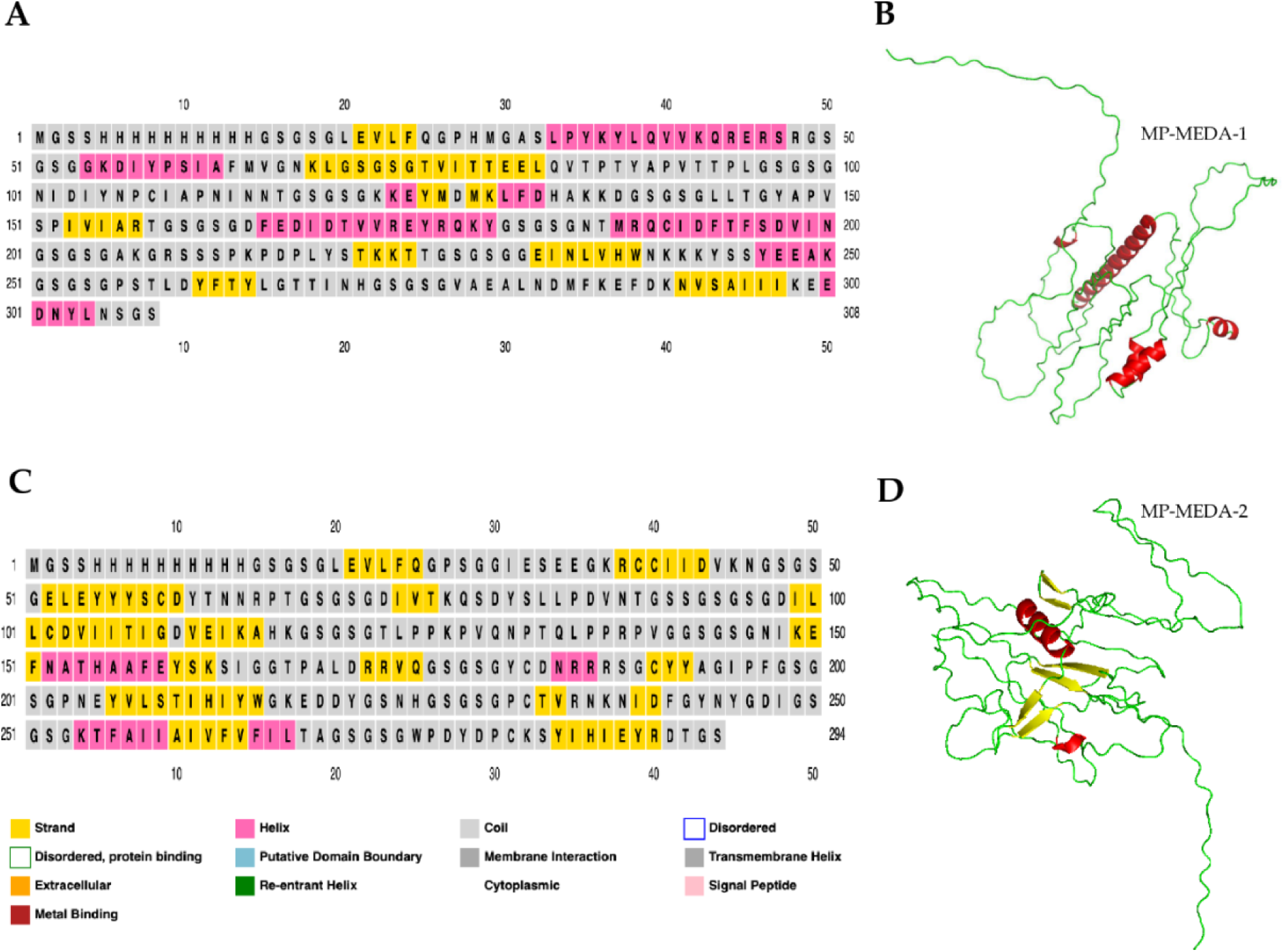
Secondary structure features of the designed diagnostic antigens. **A**. Predicted secondary structure features of the MP-MEDA-1 antigen containing 10.1% helices, 19.2% strands, and 70.7% coils. **B**. 3D structure of MP-MEDA-1, showing the different secondary structure features, including helices (red), coils (green), and strands (yellow). **C**. Predicted secondary structure features MP-MEDA-2, which contains 11.2% helices, 22.5% strands, and 66.3% coils. **D**. 3D structure of MP-MEDA-2, showing the different secondary structure features, including helices (red), coils (green), and strands (yellow).

### 3.7 Prediction, Refinement, and Validation of Modeled 3D Structures

The functional 3D structures of both chimeric antigens were predicted by AlphaFold2 on the ColabFold interface. The predicted 3D structures were subjected to two rounds of refinement, first on the FG-MD server and then on the GalaxyRefine server. The GalaxyRefine server yielded five models for each antigen. From the five predicted models, the best model for each antigen was selected for further characterization based on the obtained results. For MP-MEDA-1 (Fig. 3A), the selected model following refinement (model 2) (Fig. 3B) had the following characteristics: GDT-HA (0.8679), RMSD (0.682), and MolProbity (1.072). Additionally, the clash score was 1.6, the poor rotamer score was 0.0, and the Ramachandran plot score was 97.1%.

**Figure 3.**
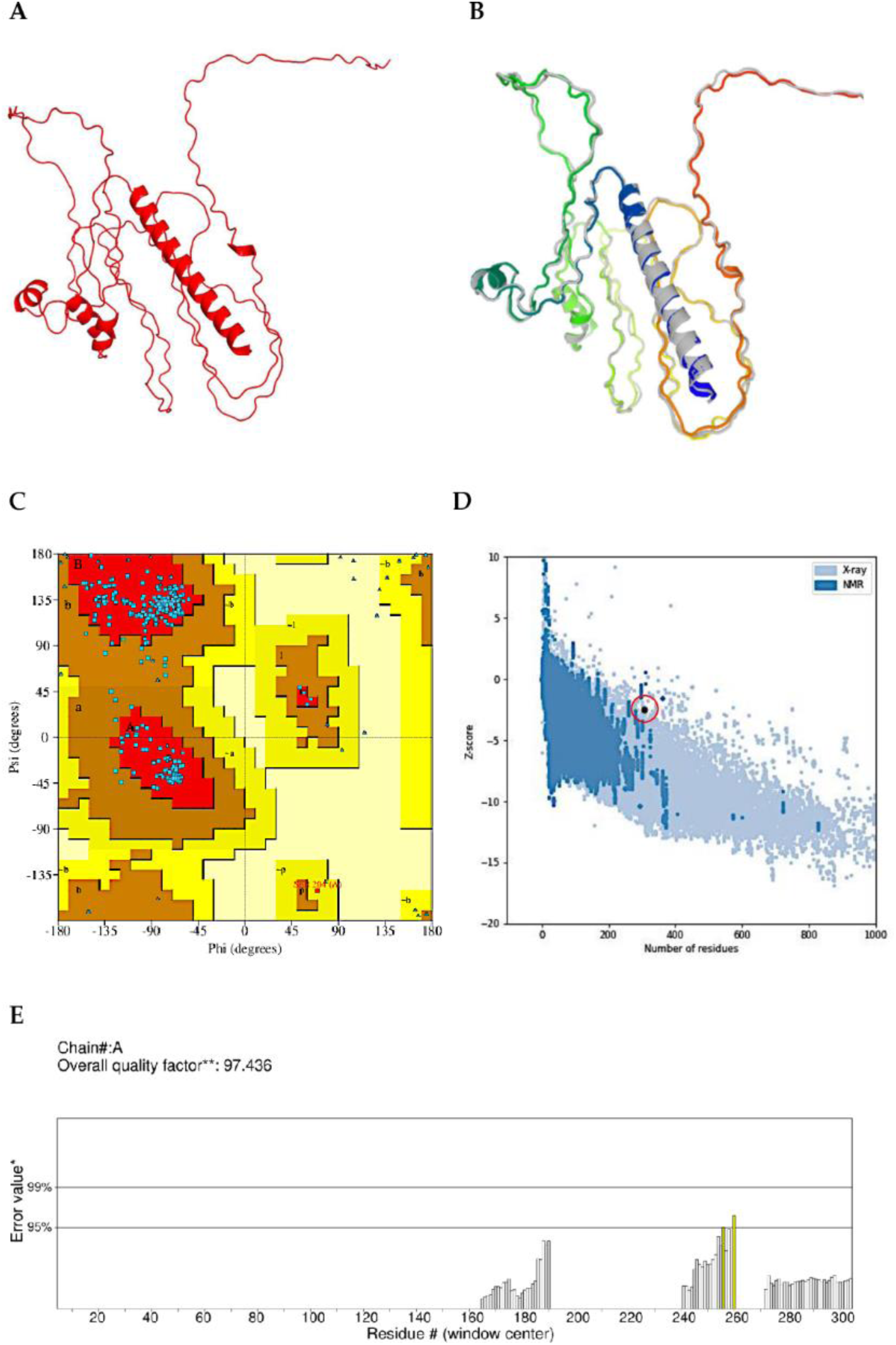
3D structure, refinement, and validation of MP-MEDA-1. **A**. The 3D structure of MP-MEDA-1 predicted by AlphaFold on the ColabFold platform. **B**. 3D structure of MP-MEDA-1 following refinement on the GalaxyRefine server. The refined 3D structure (colored) is superimposed on the initial “crude model” (gray). **C**. Ramachandran plot for MP-MEDA-1 generated by the PDBsum server, with 95.1% of the residues in the most favored region, 5.2% in the allowed region, and no residues in the disallowed regions. **D**. ProSA-web plot showing the location of MP-MEDA-1 (highlighted), with a Z score of -2.49. **E**. ERRAT quality plot for MP-MEDA-1 with a quality factor of 97.4.

For MP-MEDA-2 (Fig. 4A), the selected model (model 3) (Fig. 4B) had the following characteristics: GDT-HA (0.8333), RMSD (0.704), and MolProbity (0.949). The modeled MP-MEDA-2 structure had a clash score of 1.9, the poor rotamer score was 0.4, and the Ramachandran plot score was 98.6%. The Ramachandran plot scores predicted on the GalaxyRefine server were similar to the percentage of residues predicted to be located in favored regions by the PDBsum server: 95.1% for both MP-MEDA-1 and MP-MEDA-2. Furthermore, on basis of the results from the PDBsum server, the refined 3D structures for both MP-MEDA-1 and MP-MEDA-2 were predicted to have 4.9% of residues located in allowed regions, and 0.0% of the residues were predicted to be found in disallowed regions (Fig.3C and Fig. 4C). After 3D structure refinement, the quality of both structures was verified using the ProSA-web and ERRAT servers. The ProSA-web server predicted a Z-score of −2.49 for MP-MEDA-1 (Fig. 3D) and -2.66 for MP-MEDA-2 (Fig. 4D). On the ERRAT server, the predicted overall quality factors were 97.4 for MP-MEDA-1 (Fig. 3E) and 80.0 for MP-MEDA-2 (Fig. 4E).

**Figure 4.**
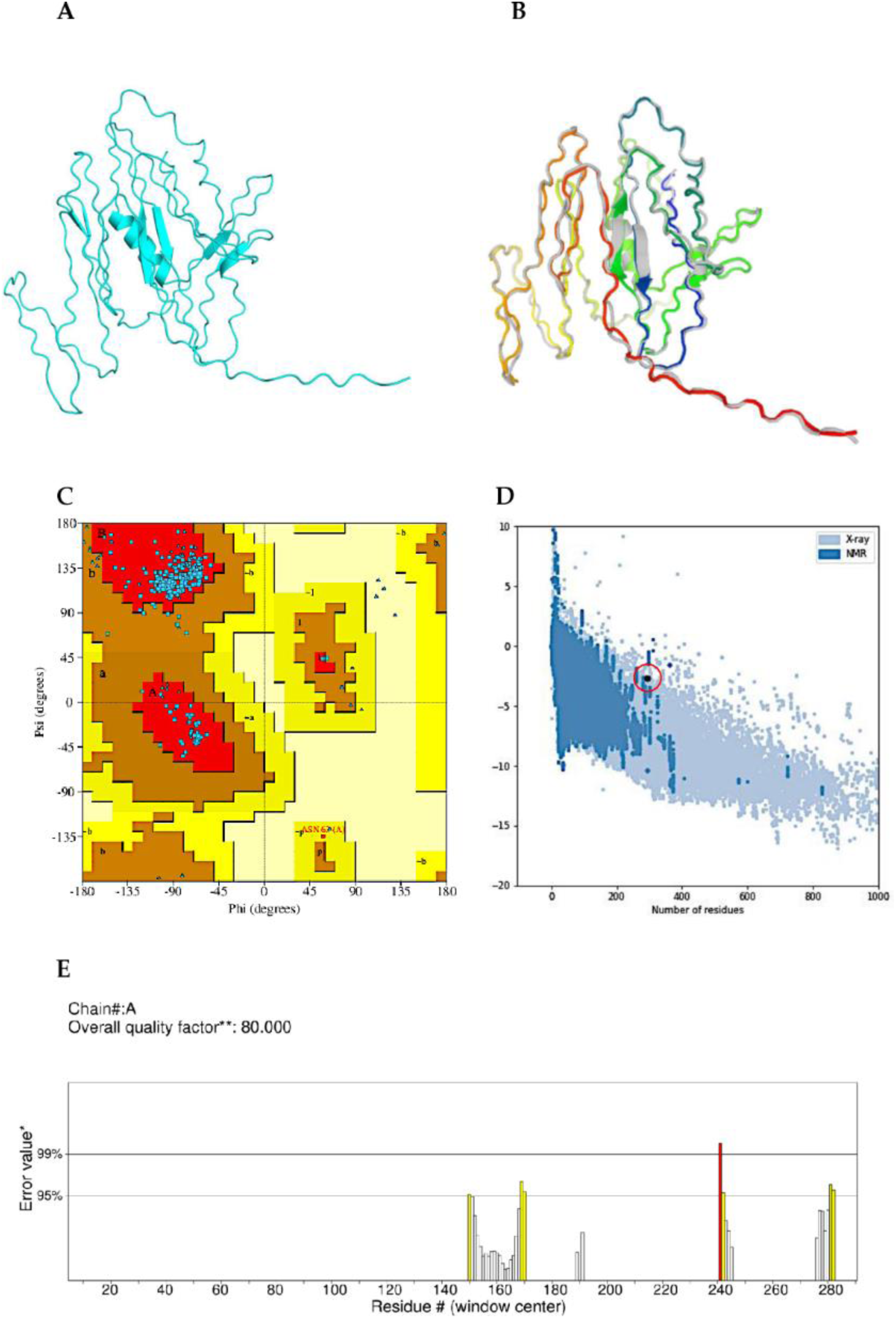
3D structure, refinement, and validation of MP-MEDA-2. **A**. 3D structure of MP-MEDA-2 predicted by AlphaFold on the ColabFold platform. **B**. The 3D structure of MP-MEDA-2 following refinement on the GalaxyRefine server. The refined 3D structure (colored) is superimposed on the initial “crude model” (gray). **C**. Ramachandran plot for MP-MEDA-2 generated by the PDBsum server, with 95.1% of the residues in the most favored region, 5.2% in the allowed region, and no residues in the disallowed regions. **D**. ProSA-web plot showing the location of MP-MEDA-2 (highlighted), with a Z score of -2.66. **E**. ERRAT quality plot for MP-MEDA-2 with a quality factor of 80.0.

### 3.8 Discontinuous B-cell epitope prediction

This was performed using the ElliPro server. For MP-MEDA-1, a total of one hundred and forty-nine amino acid residues (48.4%) were predicted to be found in 16 conformational B-cell epitopes. The number of residues in the predicted epitopes ranged from 4 to 18, and the epitope prediction scores ranged from 0.587 - 0.968 (Table S2). For MP-MEDA-2, on the other hand, a total of one hundred and fifty-two amino acid residues (51.7%) were predicted to be located in 15 conformational B-cell epitopes. The number of residues in the predicted epitopes ranged from 3 to 20, and the epitope prediction scores ranged from 0.522 - 0.980 (Table S3).

### 3.9 Protein–protein docking between the designed diagnostic antigens and immunoglobulins

Suitable immune interactions between antigens and antibodies are required to develop serological tests such as ELISA (101). Several assays have been developed for the serological testing of Mpox using IgA, IgG, and IgM (25, 26, 102, 103). Protein‒protein docking on the Cluspro 2.0 server was performed to investigate the binding interaction between the validated 3D structures of the multiepitope diagnostic antigens and the Fab regions of the selected immunoglobulins. From the thirty models generated, displayed, and ranked by cluster size on the ClusPro 2.0 server, the docked complex with the largest cluster size for each interaction was selected. For the MP-MEDA-1-Ig complexes, the ClusPro 2.0 results predicted that the highest number of docking poses clustered had the lowest energy-weighted scores of -342.6, - 342.6, and -360.1 for IgA, IgG, and IgM, respectively (Fig. 5A, C, and E). For the MP-MEDA-2-Ig complexes, ClusPro 2.0 results predicted that the highest number of docking poses clustered had the lowest energy-weighted scores of -390.7, -379.8, and -375.7 for IgA, IgG, and IgM, respectively (Fig. 5B, D, and F). For further characterization of diagnostic antigen– Ig Fab interactions, the complexes formed by the interactions of MP-MEDA-1 and MP-MEDA-2 with the selected Ig Fab regions were further analyzed for their binding affinities using the PrODIGY and AREA-AFFINITY servers. In addition, the interactions at the interface of the formed complexes were analyzed on the PDBsum server. For MP-MEDA-1, the predicted relative binding free energies (ΔG) of the antigen–Ig Fab complexes at 25 °C for IgA, IgG, and IgM were -14.1 kcal/mol, -11.2 kcal/mol, and -14.8 kcal/mol, respectively. Furthermore, the predicted (dissociation constant) K_d_ values for the interactions with IgA, IgG, and IgM were 4.5 × 10^−11^, 6.6 × 10^−9^, and 1.3 × 10^−11^, respectively. Additionally, the number of contacts formed at the interface (IC) per property was determined as follows: (ICs charged– charged: 9, ICs charged–polar: 21, ICs charged–apolar: 23, ICs polar–polar: 14, ICs polar– apolar: 35, and ICs apolar–apolar: 10 for IgA. For the IgG, the number of contacts formed at the interface (IC) per property was determined as follows: (ICs charged–charged: 6, ICs charged–polar: 18, ICs charged–apolar: 22, ICs polar–polar: 14, ICs polar–apolar: 24 and ICs apolar–apolar: 15. Moreover, for IgM, the number of contacts formed at the interface (IC) per property was determined as follows: (ICs charged–charged: 5, ICs charged–polar: 25, ICs charged–apolar: 27, ICs polar–polar: 16, ICs polar–apolar: 40 and ICs apolar–apolar: 16 (Table 4).

**Figure 5.**
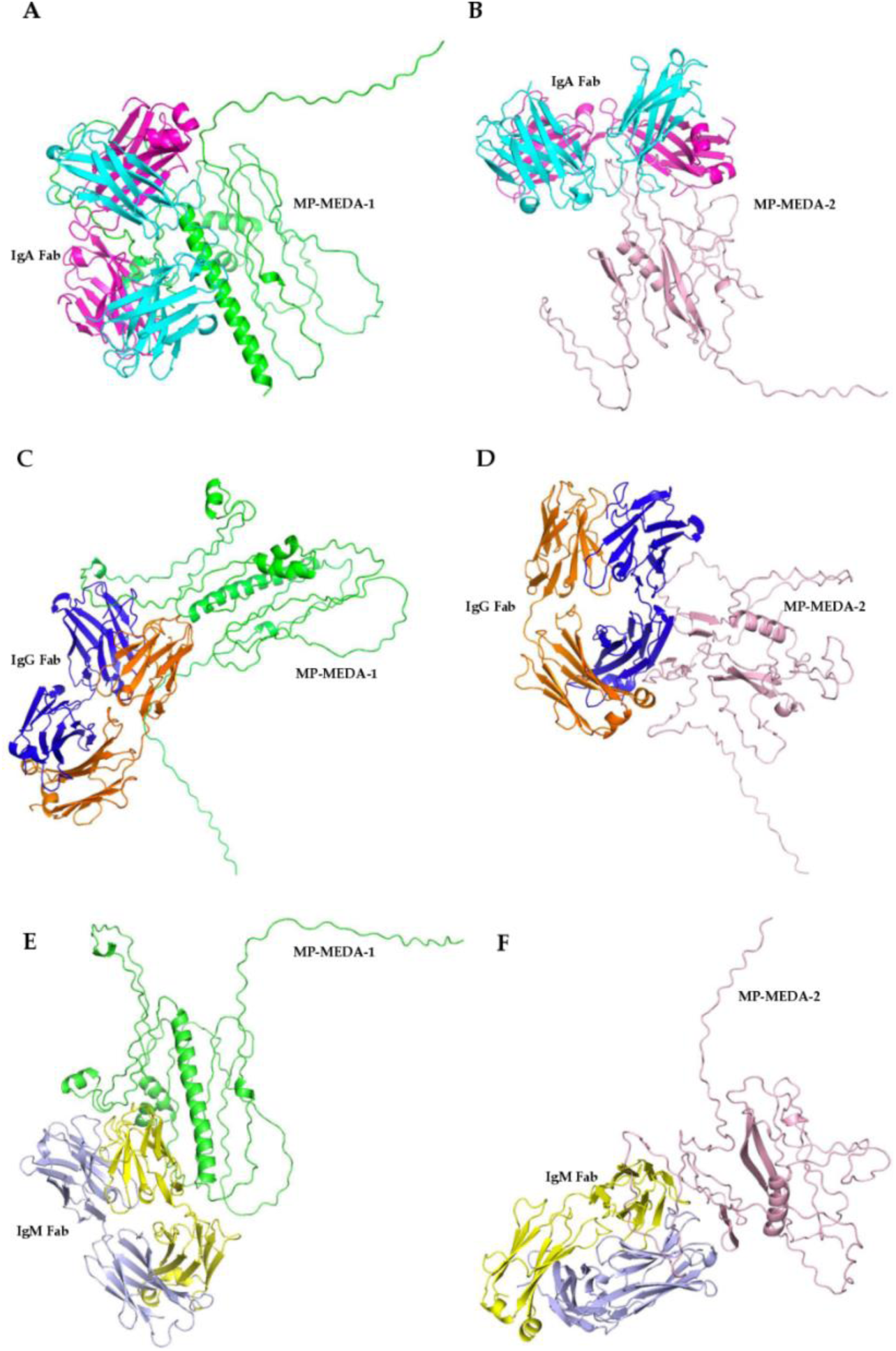
Interactions of designed multiepitope diagnostic antigens with Ig Fab regions. **A**. Structure of the complex formed between MP-MEDA-1 (green) and the IgA Fab region. The heavy chain is in magenta, whereas the light chain is in pink. **B**. Structure of the complex formed between MP-MEDA-2 (light pink) and the IgA Fab region. The heavy chain is in magenta, whereas the light chain is in pink. **C**. Structure of the complex formed between MP-MEDA-1 (green) and the IgG Fab region. The heavy chain is in blue, whereas the light chain is in orange. **D**. Structure of the complexes formed between MP-MEDA-2 (light pink) and the IgG Fab region. The heavy chain is blue while the light chain is orange. **E**. Structure of the complex formed between MP-MEDA-1 (green) and the IgM Fab region. The heavy chain is light purple, while the light chain is in yellow. F. Structure of the complex formed between MP-MEDA-2 (light pink) and the IgM Fab region. The heavy chain is light purple, while the light chain is yellow.

**Table 4:** Binding interactions with the designed diagnostic antigens and the Fab regions of IgA, IgG, and IgM.

| Protein-protein<br>complex | $\Delta G$<br>(kcal<br>mol<sup>-1</sup>) | $K_d$ (M)<br>at 25<br>°C | ICs<br>charged-<br>charged | ICs<br>charged-<br>polar | ICs<br>charged-<br>apolar | ICs<br>polar-<br>polar | ICs<br>polar-<br>apolar | ICs<br>apolar-<br>apolar | NIS<br>charged | NIS<br>apolar |
| --- | --- | --- | --- | --- | --- | --- | --- | --- | --- | --- |
| IgA/MP-MEDA-1 | -14.1 | 4.5e-11 | 9 | 21 | 23 | 14 | 35 | 10 | 20.11 | 39.67 |
| IgG/MP-MEDA-1 | -11.2 | 6.6e-09 | 6 | 18 | 22 | 14 | 24 | 15 | 19.17 | 40.73 |
| IgM/MP-MEDA-1 | -14.8 | 1.3e-11 | 5 | 25 | 27 | 16 | 40 | 16 | 20.97 | 39.19 |
| IgA/MP-MEDA-2 | -17.7 | 1e-13 | 4 | 23 | 23 | 20 | 58 | 36 | 18.96 | 40.27 |
| IgG/MP-MEDA-2 | -14.2 | 3.6e-11 | 9 | 26 | 28 | 9 | 29 | 13 | 18.42 | 40.79 |
| IgM/MP-MEDA-2 | -10.3 | 2.9e-08 | 8 | 9 | 22 | 4 | 10 | 13 | 20.07 | 39.31 |

For MP-MEDA-2, the predicted relative binding free energies (ΔG) of the antigen–Ig Fab complexes at 25 °C for IgA, IgG, and IgM were -17.7 kcal/mol, -14.2 kcal/mol, and -10.3 kcal/mol, respectively. In addition, the predicted K_d_ values for the interactions with IgA, IgG, and IgM were 1.0 × 10^−13^, 3.6 × 10^−11^, and 2.9 × 10^−8^, respectively (Table 4). In addition, the number of contacts formed at the interface (IC) per property was determined as follows: (ICs charged–charged: 4, ICs charged–polar: 23, ICs charged–apolar: 23, ICs polar–polar: 20, ICs polar–apolar: 58, and ICs apolar–apolar: 36 for IgA. For the IgG, the number of contacts formed at the interface (IC) per property was determined as follows: (ICs charged–charged: 9, ICs charged–polar: 26, ICs charged–apolar: 28, ICs polar–polar: 9, ICs polar–apolar: 29 and ICs apolar–apolar: 13. Moreover, for IgM, the number of contacts formed at the interface (IC) per property was determined as follows: (ICs charged–charged: 8, ICs charged–polar: 9, ICs charged–apolar: 22, ICs polar–polar: 4, ICs polar–apolar: 10 and ICs apolar–apolar: 13 (Table 4). In addition, all the antigen-Ig Fab complexes were predicted to comprise several clusters with multiple residues from the antigen interacting with the antibody heavy and light chains (Table S5)

Coupled with the binding interaction data obtained from the Cluspro 2.0 server, the binding affinity and binding energy were also predicted via the AREA-AFFINITY web server. For MP-MEDA-1, the predicted binding energies of the antigen–Ig Fab complexes were -10.5358347 kcal/mol, -9.7613837 kcal/mol, and -12.5425784 kcal/mol for IgA, IgG, and IgM, respectively. In addition, the predicted binding affinities (log(K)) for the interactions between the antigen and IgA, IgG, and IgM were -7.7247853, -7.1569643, and -9.1961129, respectively. Moreover, for MP-MEDA-2, the predicted binding energies of the antigen–Ig Fab complexes were - 10.5358347 kcal/mol, -9.7613837 kcal/mol, and -12.5425784 kcal/mol for IgA, IgG, and IgM, respectively. In addition, the predicted binding affinities (log(K)) for the interactions between the antigen and IgA, IgG, and IgM were -7.7247853, -7.1569643, and -9.1961129, respectively (Table 5). In summary, both diagnostic antigens exhibited favourable interactions with the selected immunoglobulins (IgA, IgG, and IgM).

**Table 5:** Predicted affinities and binding energies between MP-MEDA-1 and MP-MEDA-2 and the Fab regions of IgA, IgG, and IgM.

| Antibody | MP-MEDA-1 |  | MP-MEDA-2 |  |
| --- | --- | --- | --- | --- |
|  | Predicted binding affinity (log(K)) | Predicted binding energy (kcal/mol) | Predicted binding affinity (log(K)) | Predicted binding energy (kcal/mol) |
| **IgA** | -7.7247853 | -10.5358347 | -8.7184647 | -11.8911140 |
| **IgG** | -7.1569643 | -9.7613837 | -7.9518258 | -10.8454951 |
| **IgM** | -9.1961129 | -12.5425784 | -8.3071034 | -11.3300584 |

Interaction analyses on the PDBsum server revealed that the designed antigens have multiple interactions with the heavy and light chains of the Fab regions of IgA, IgG, and IgM. The interaction analyses included the number of residues at the interfaces of each complex, the area of the interface, the number of salt bridges, disulfide bridges, H-bonds, and non-bonded contacts. The IgA/MP-MEDA-1 complex presented the highest number of interface residues between the designed antigen and the Ig Fab region. This complex also had the highest number of non-bonded contacts. The least number of interface residues, as well as non-bonded contacts, was predicted for the IgM/MP-MEDA-2. All of the complexes were predicted to contain at least one salt bridge and 10-H bonds between the antigen and the selected Ig Fab region. Disulfide bridges were predicted only between the heavy and light chains of the Ig Fab regions (Table S4). In addition, the data from the PICKLUSTER tool revealed multiple residues in different clusters at the interfaces between the designed antigens and the heavy and light chains of the Fab regions in the complexes. The cluster with the greatest number of interacting residues was predicted between MEDA-1 and the heavy chain of the Fab region of IgA (Table S5).

### 3.10 Molecular dynamics simulation

Following protein‒protein docking, molecular dynamics simulation was performed by the iMODs server to investigate the stability of the complexes formed by the interactions between the designed diagnostic antigens and the Fab regions of IgA, IgG, and IgM. A significant degree of distortion was observed in all the complexes, with each residue in the complex playing a role in their stability (Figs. 6A, 7A, 8A, 9A, 10A, and 11A). The B-factor plots (Fig. 6B, 7B, 8B, 9B, 10B, and 11B) indicate the correlation between the mobility of the docked complex NMA and the PDB score (proportional to the mean RSMD). The predicted eigenvalues for the interactions between MP-MEDA-1 and the IgA, IgG, and IgM Fab regions were 5.33247 × 10^−7^, 8.210414 × 10^−6^, and 5.940199 × 10^−6^, respectively (Figs. 6C, 7C, and 8C). For MP-MEDA-2, the predicted eigenvalues were 1.028857 × 10^−5^, 8.0122801 × 10^−6^, and 7.373463 × 10^−6^ for the interactions with the IgA, IgG, and IgM Fab regions, respectively (Figs. 9C, 10C, and 11C). These values indicate the energy required to deform the structure. A lower value means easier deformation. Each normal mode of the formed complexes is linked to variance plots that represent both the individual variances (in purple) and the cumulative variances (in green) (Figs. 6D, 7D, 8D, 9D, 10D, and 11D). The elastic network maps predicted the intricate connections between pairs of atoms in all the different complexes. The grey dots indicate stiffer regions, whereas the lighter dots indicate more flexible regions (Figs. 6E, 7E, 8E, 9E, 10E, and 11E). In addition, the covariance matrix for each of the complexes revealed the patterns between pairs of residues in the dynamic regions of the complexes (related, unrelated, and anti-related motions in complex structures are represented in blue, red, and white, respectively) (Fig. 6F, 7F, 8F, 9F, 10F, and 11F). These results suggest that the designed multiepitope diagnostics antigens can interact with the Fab regions of IgA, IgG, and IgM, forming complexes with favourable molecular stability. The results from the AGGRESCAN3D v.2.0 server predicted that all complexes formed between the designed antigens and the selected Ig Fab regions contained significant proportions of soluble and aggregation-prone residues (Fig. S3).

**Figure 6.**
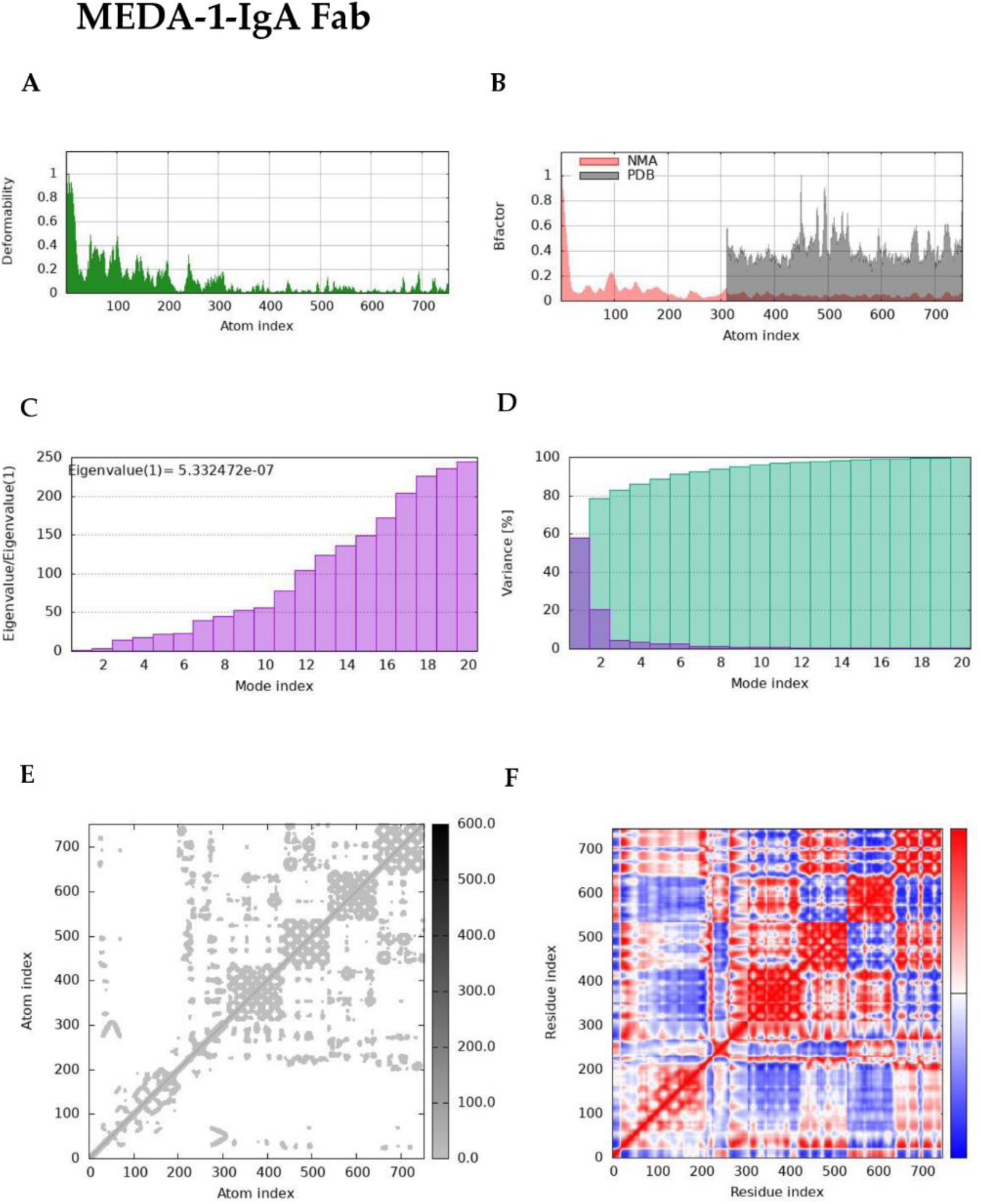
Molecular dynamics simulation of the complex between MP-MEDA-1 and the IgA Fab region. Molecular dynamics simulation predictions showing **A**. deformability. **B**. B-factor **C**. Eigenvalues (a lower value indicates easier deformation). **D**. Variance (purple bars: individual variances and green bars: cumulative variances). **E**. Elastic network map (darker regions indicate stiffer regions) and **F**. Covariance map (red: correlated, white: uncorrelated, blue: anti-correlated).

**Figure 7.**
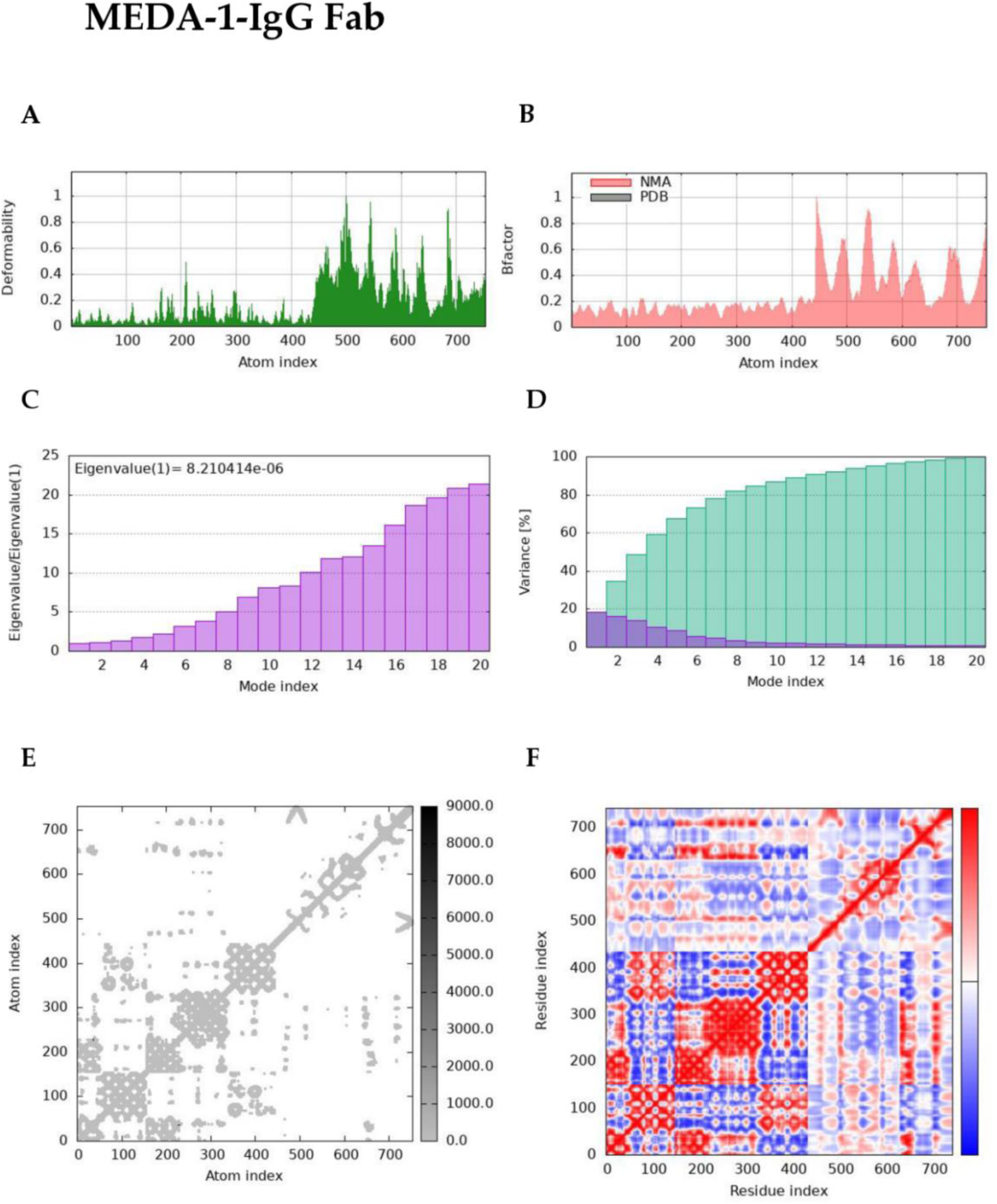
Molecular dynamics simulation of the complex between MP-MEDA-1 and the IgG Fab region. Molecular dynamics simulation predictions showing **A**. deformability. **B**. B-factor **C**. Eigenvalues (a lower value indicates easier deformation). **D**. Variance (purple bars: individual variances and green bars: cumulative variances). **E**. Elastic network map (darker regions indicate stiffer regions) and **F**. Covariance map (red: correlated, white: uncorrelated, blue: anti-correlated).

**Figure 8.**
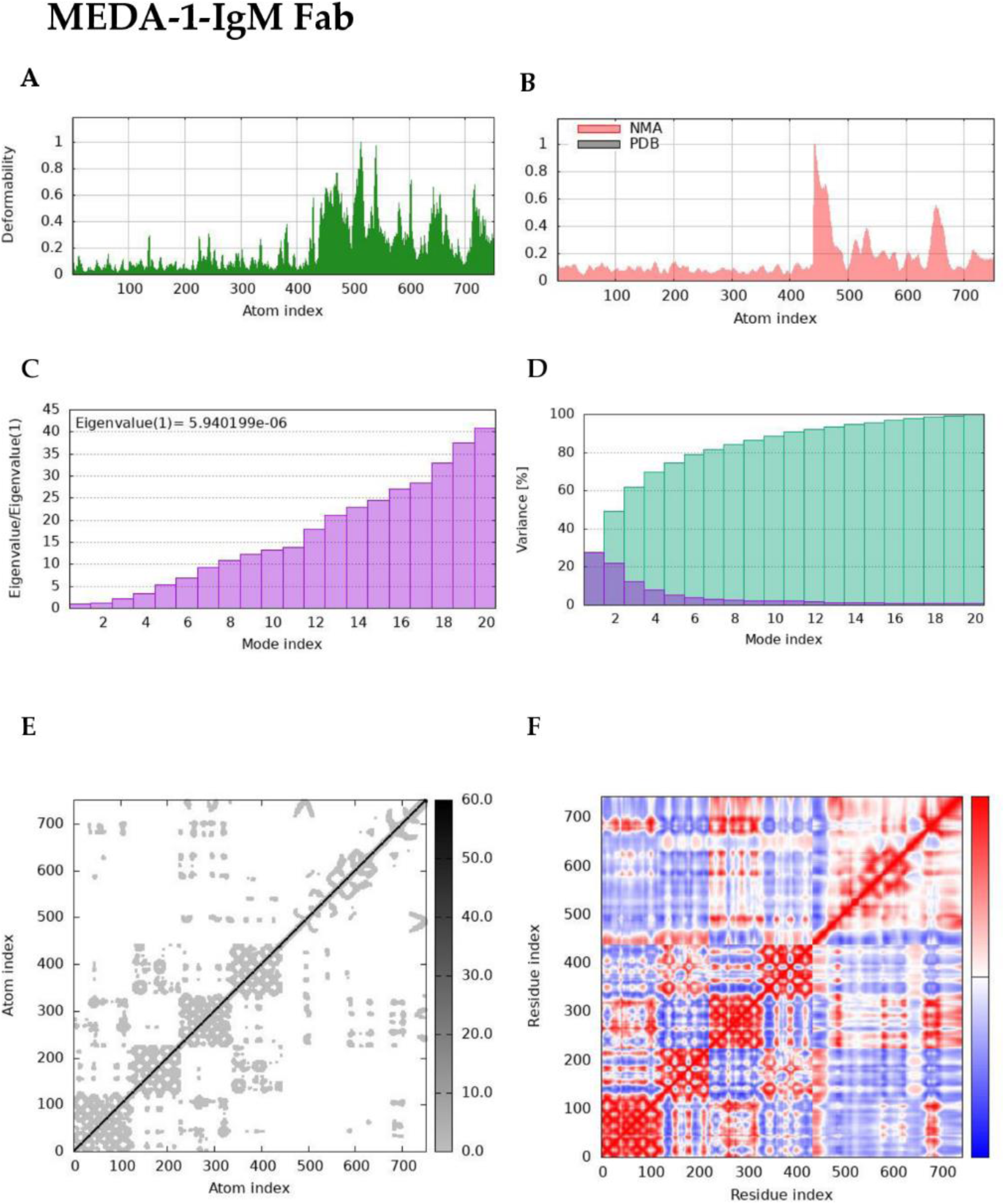
Molecular dynamics simulation of the complex between MP-MEDA-1 and the IgM Fab region. Molecular dynamics simulation predictions showing **A**. deformability. **B**. B-factor **C**. Eigenvalues (a lower value indicates easier deformation). **D**. Variance (purple bars: individual variances and green bars: cumulative variances). **E**. Elastic network map (darker regions indicate stiffer regions) and **F**. Covariance map (red: correlated, white: uncorrelated, blue: anti-correlated).

**Figure 9.**
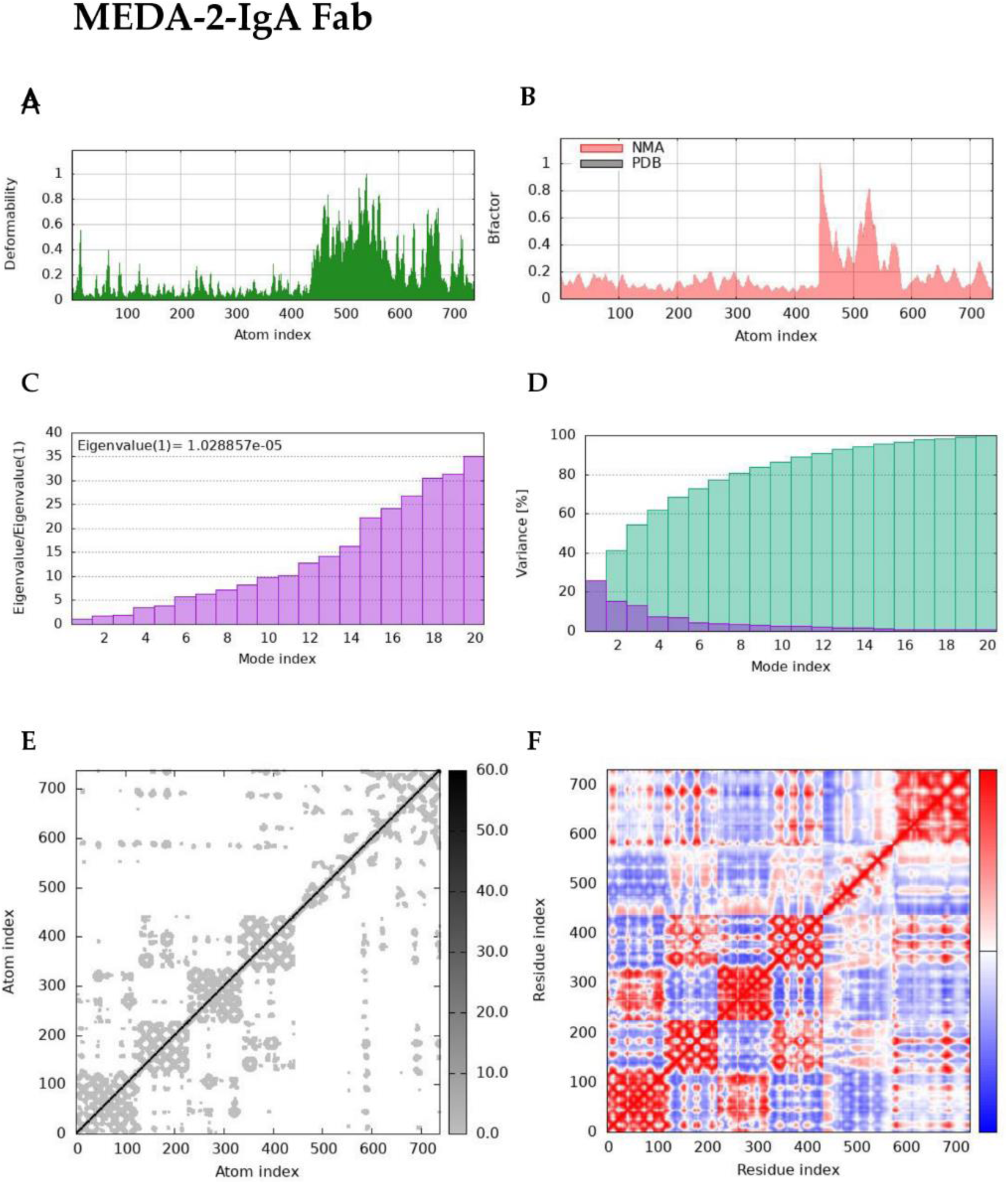
Molecular dynamics simulation of the complex between MP-MEDA-2 and the IgA Fab region. Molecular dynamics simulation predictions showing **A**. deformability. **B**. B-factor **C**. Eigenvalues (a lower value indicates easier deformation). **D**. Variance (purple bars: individual variances and green bars: cumulative variances). **E**. Elastic network map (darker regions indicate stiffer regions) and **F**. Covariance map (red: correlated, white: uncorrelated, blue: anti-correlated).

**Figure 10.**
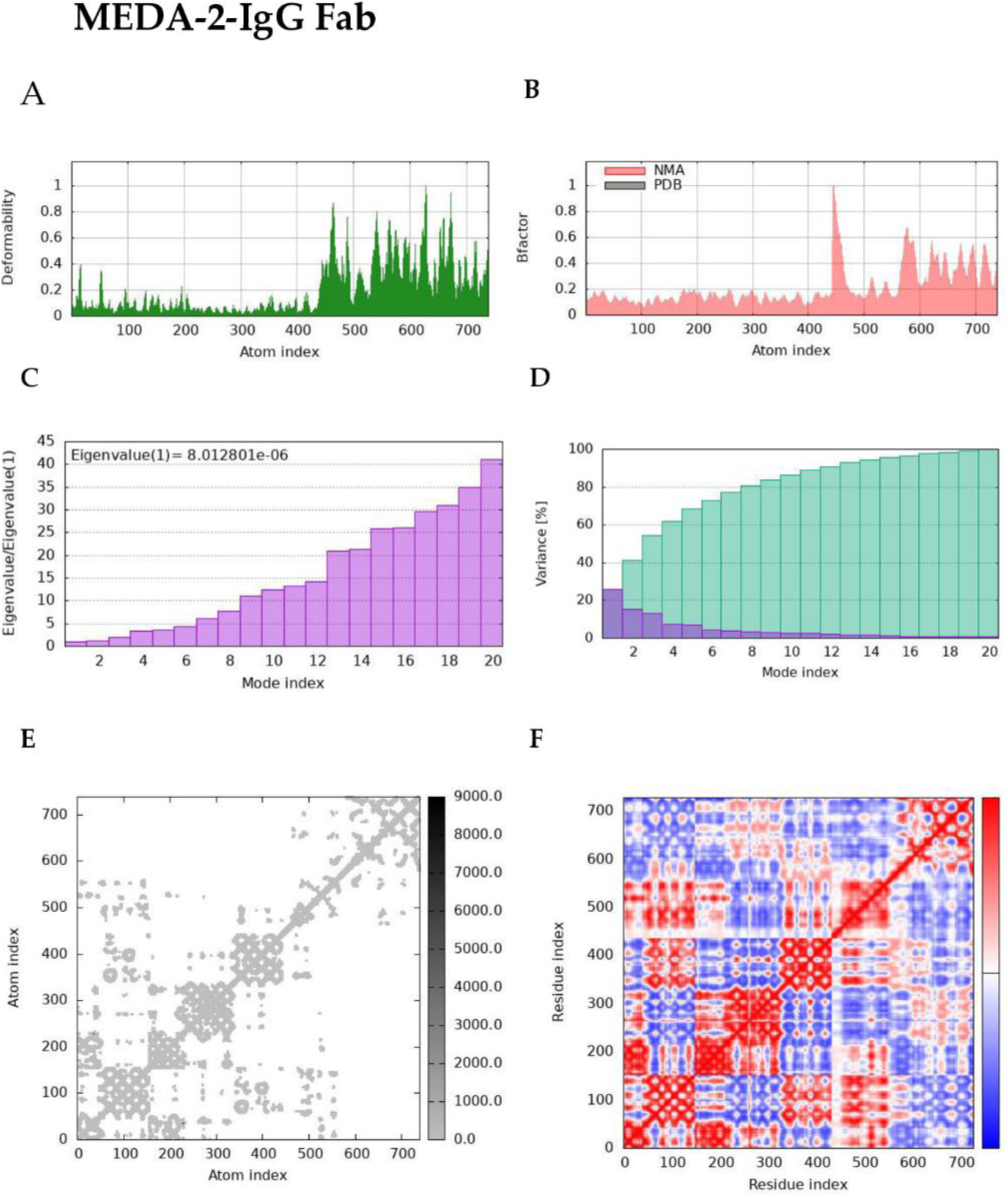
Molecular dynamics simulation of the complex between MP-MEDA-2 and the IgG Fab region. Molecular dynamics simulation predictions showing **A**. deformability. **B**. B-factor **C**. Eigenvalues (a lower value indicates easier deformation). **D**. Variance (purple bars: individual variances and green bars: cumulative variances). **E**. Elastic network map (darker regions indicate stiffer regions) and **F**. Covariance map (red: correlated, white: uncorrelated, blue: anti-correlated).

**Figure 11.**
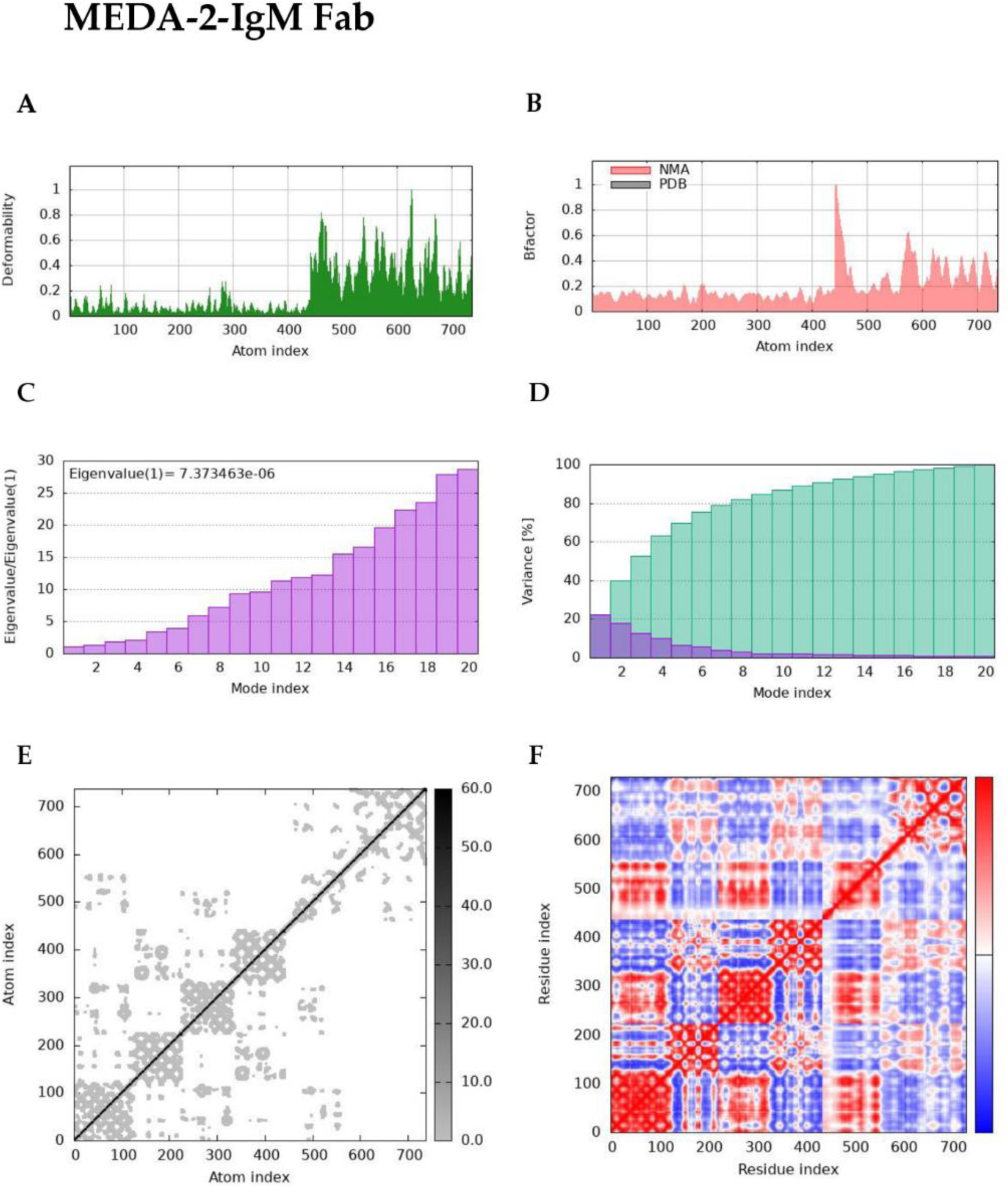
Molecular dynamics simulation of the complex between MP-MEDA-2 and the IgM Fab region. Molecular dynamics simulation predictions showing **A**. deformability. **B**. B-factor **C**. Eigenvalues (a lower value indicates easier deformation). **D**. Variance (purple bars: individual variances and green bars: cumulative variances). **E**. Elastic network map (darker regions indicate stiffer regions) and **F**. Covariance map (red: correlated, white: uncorrelated, blue: anti-correlated).

### 3.11 Gene Sequence Codon Optimization and In Silico Restriction Cloning

The Java Codon Adaptation Tool (JCat) server was used to perform codon optimization on the gene sequences encoding for the designed multiepitope diagnostic antigens for expression in *the E. coli* K12 strain. The codon-optimized sequences contained 924 nucleotides for MP-MEDA-1 and 882 nucleotides for MP-MEDA-2. A predicted codon adaptation index (CAI) of 1.0 was predicted for both sequences, whereas GC contents of 49.2% and 51.7% were predicted for MP-MEDA-1 and MP-MEDA-2, respectively. The GC contents of the codon-optimized sequences were close to the 50.7% GC content of the *E. coli* K12 strain, suggesting that both proteins can be suitably expressed in bacteria. Generally, the GC content of bacterial genomes ranges from 25% to 75% (104), and a GC content between 30% and 70% is recommended for optimal protein expression (105). In addition, the predicted CAI for each codon-optimized sequence suggests the possibility of high-level expression in bacteria. Lastly, in-silico cloning was performed to insert the codon-optimized sequences for both antigens into the bacterial vector pET30a (+) via SnapGene software v8.0.2 (Fig. 12).

**Figure 12.**
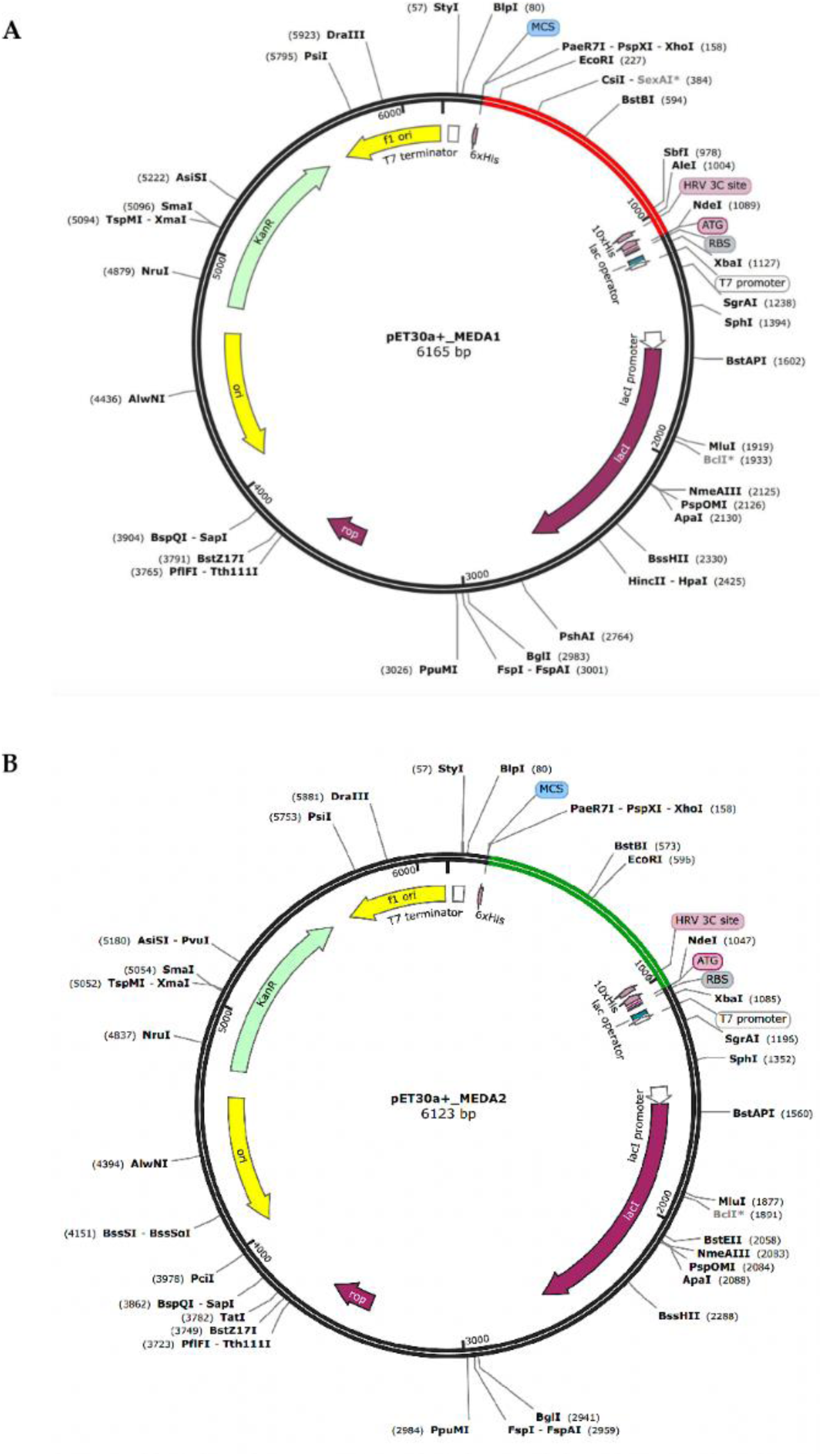
In silico cloning of codon-optimized DNA sequences coding for the designed chimeric antigens. The codon-optimized sequences were cloned between the NdeI and XhoI sites of the pET30a+ vector for expression in *the E. coli* K12 strain. (**A**) The inserted DNA sequence is shown in red for MP-MEDA-1 and green for MP-MEDA-2 (**B**). The plasmid backbone is in black. The 10×His tags and the HRV 3C protease cleavage sites are highlighted in both insert sequences.

### 3.12 RNA secondary and tertiary structure prediction

The RNAfold server was used to predict the secondary structures of the mRNA sequences coding for both designed antigens. The thermostability of both mRNA secondary sequences was demonstrated by the predicted negative free energies for both antigens. For MP-MEDA-1, the analysis predicted a minimal free energy (MFE) of -264.00 kcal/mol for the optimal secondary structure (Fig. 13A, C) and MFE of the centroid structure of -194.22 kcal/mol (Fig. S4A). For MP-MEDA-2, the predicted MFE for the optimal secondary structure was -268.60 kcal/mol (Fig. 13B, D), while the MFE of the centroid structure was -195.12 kcal/mol (Fig. S4B). In addition, the free energies of the thermodynamic ensembles were -278.62 kcal/mol for MP-MEDA-1 and -280.75 kcal/mol for MP-MEDA-2. The tertiary structures of the transcribed mRNA sequences were also predicted using the trRosettaRNA server. Both predicted tertiary structures, however, had pLDDTs of less than 50, indicating a low confidence in the structure (Fig. S5).

**Figure 13.**
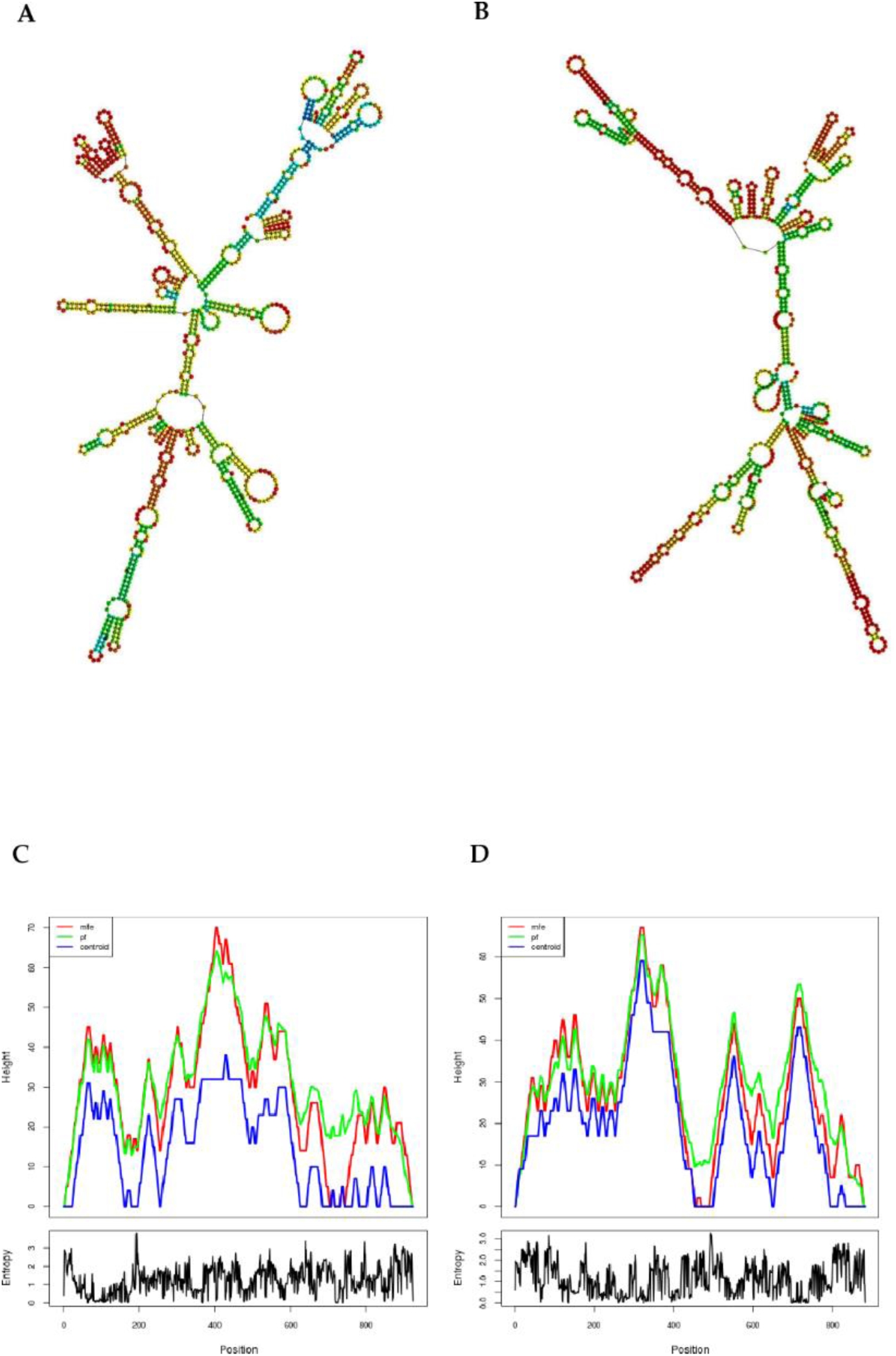
Predicted secondary structures of the MP-MEDA-1 and MP-MEDA-2 mRNA sequences. The MFE secondary structures of the mRNA sequences for MP-MEDA-1 (A) and MP-MEDA-2 (**B**). The mountain plots for the mRNA secondary structures indicate the minimal free energy of the mRNA secondary structures for MP-MEDA-1 (**C**) and MP-MEDA-2 (**D**).

## 4 Discussion

Given the public health and socioeconomic impact of the last Mpox outbreaks, there has been a consensus that continuous surveillance will be imperative in strengthening the response and mitigating the impact of the outbreak, especially in sub-Saharan Africa, where the highest numbers of cases have been reported (106). Although genomic surveillance can generate very valuable data, many African countries lack the capacity to execute large-scale genomic surveillance studies, and it is likely that the virus is circulating but remains undetected owing to limited genomic surveillance (107, 108). Serosurveillance studies can provide valuable data for understanding population immunity, tracking viral transmission, and guiding public health decision-making (109). A serological assay to assess immunity to Mpox is urgently needed to enhance the current understanding of humoral immune responses to both Mpox infection and vaccination (110). Multiepitope antigens based on B-cell epitopes have been proposed as an approach to serodiagnostic test development with increased sensitivity and enhanced specificity (111, 112) and have been tested for several diseases. This project focused on the use of computational strategies to design and characterize two chimeric multiepitope antigens for Mpox serosurveillance (Fig. S6).

To design multiepitope diagnostic antigens that can be used for Mpox serosurveillance, five viral proteins previously implicated in the development of diagnostic tools were used to predict linear B-cell epitopes, and antigenic epitopes were selected to design two antigens (MP-MEDA-1 and MP-MEDA-2). The designed chimeric antigens were predicted to be antigenic, with MP-MEDA-2 predicted to be slightly more antigenic than MP-MEDA-1, based on results from the VaxiJen v2.0 and ANTIGENpro servers (Table 2). The antigenicity of both diagnostic antigens supports their potential deployment as tools for Mpox serosurveillance since antigenicity refers to an interaction between antigenic determinants (epitopes) and antibodies or specific T-cell receptors for an antigen (113). The antigenicity of the designed chimeric antigens was also supported by the presence of large proportions of exposed residues (and interacting residues), which could interact with antibodies in the development of antibody-capture tests for serosurveillance (114).

The designed diagnostic antigens were predicted to possess suitable physicochemical properties. The predicted instability index revealed that although MP-MEDA-1 was more stable, MP-MEDA-2 was slightly unstable since the II (42.32) was still close to the threshold of 40. On the other hand, the GRAVY values and aliphatic index values predicted for both antigens indicated that both diagnostic antigens were hydrophilic and exhibited thermal stability. The hydrophilic nature of both proteins is supported by the large proportions of hydrophilic amino acids present in the sequences (Table 3). In addition, the presence of these hydrophilic amino acids could enhance the stability of antigen‒antibody complexes through charge‒charge interactions (115). The physicochemical properties of both designed antigens support their role as tools for Mpox serosurveillance.

The 3D structures of both antigens were predicted, refined, and validated via the ProsA-web and ERRAT servers. The Z-scores of the validated 3D structures fell within the range for proteins of comparable sizes predicted through X-ray crystallography. In addition, the Ramachandran and ERRAT plots suggested a high confidence in the predicted 3D structures. For Ramachandran plots, a high percentage of residues in the most favored regions (> 90%) is generally regarded as a good indicator of high-quality protein structure (116). The predicted ERRAT scores also suggested the high quality of the 3D structures of both proteins. ERRAT scores of 80 and above generally suggest accurate predictions (117). Since antigen-specific IgA, IgG, and IgM antibodies have been detected in Mpox-infected patients (26, 27), the Fab regions of selected immunoglobulins were used to investigate the interactions with the designed antigens. The Fab regions were selected for convenience since they are responsible for antigen recognition through their variable sites (118). Binding energy, binding affinity, and interaction analyses suggested that MP-MEDA-1 and MP-MEDA-2 formed stable complexes with the Fab region of all the selected immunoglobulins (IgA, IgG, and IgM), with several residues at the antigen‒antibody complex interfaces. From the PRODIGY server, for MP-MEDA-1, the lowest ΔG was predicted for the interaction with the Fab region of IgM. In contrast, the lowest K_d_ (dissociation constant) was predicted for the interaction with the Fab region of IgA. However, the ΔGs for the interactions between IgA and IgM were very similar (Table 4). For MP-MEDA-2, the lowest ΔG and K_d_ values were predicted for the interaction with the Fab region of IgA. Nevertheless, all the complexes formed between the designed antigens and Fab regions of the selected immunoglobulins presented low K_d_ values, indicating high-affinity interactions. In ELISA, the affinity of the antigen-antibody interaction has been reported to be crucial for assay sensitivity and accuracy (119). NMA analyses were performed to analyze the molecular mobility and comparative deformability of the complexes formed between the designed antigens and the Ig Fab regions. The data obtained (including low eigenvalue values) indicated favorable flexibility of all the docked complexes.

The *E. coli* system was selected to express the multiepitope subunit diagnostic antigens since it is the most efficient host system for recombinant protein expression (120). Codon optimization, as well as the predicted MFE for the mRNA secondary structures, supports the possibility of high levels of expression for both proteins. However, mRNA structure prediction yielded structures with low confidence. This implies that the mRNA structures may not be very reliable.

A key limitation of this study is that, thus far, all the data have been generated using computational tools. It would be important for MP-MEDA-1 and MP-MEDA-2 to be expressed and assessed in serological assays to establish them as tools for Mpox serosurveillance. Nevertheless, given the success of this approach in the computational design of diagnostic antigens for other pathogens, the data obtained provide valuable insights into the potential role of the designed diagnostic candidates as serological tools for Mpox control.

## 5 Conclusions

In the last few years, two Mpox outbreaks have occurred, with the virus spreading across more than 100 countries, leading to multiple deaths and causing severe social and economic disruptions on a global scale. In the present study, two multiepitope chimeric antigens (containing linear B-cell epitopes) were designed using computational techniques. The designed multiepitope diagnostic antigens demonstrated favorable features (including antigenicity, thermal stability, and 3D structural features). In addition, the designed antigens demonstrated stable interactions with the Fab regions of IgA, IgG, and IgM (which have been reported to be involved in Mpox progression and have been assessed in serodiagnosis. From the results obtained, it can be concluded that MP-MEDA-1 and MP-MEDA-2 demonstrate features that can be further harnessed in the development of diagnostic tools for Mpox serosurveillance. Nevertheless, it is important to validate the computational results obtained from this study in vitro and in vivo experiments.

## Declarations

### Ethics approval and consent to participate

Not applicable.

### Consent for publication

Not applicable

### Availability of data and materials

All the data generated or analyzed during this study are included in this published article (and its supplementary information files).

### Authors’ contributions

RAN, RAS, SMG, JS, LV and VPKT designed the project; RAN, RAS, MTE, DNN, TYAS, ABA, NLA, BTT, KYG and GTN performed the experiments; RAN, RAS, KYG, JAC, DNN, CMS, NEY, NLA, BTT, TBM and BNY evaluated and interpreted the data; RAN, RAS, KYG, JEE, NEY, CMS, BNY, JAC, TBM and NLA prepared the draft manuscript; RAS and GTN prepared the figures; RAS, RAN, GTN, TYAS, and JEE finalized the manuscript; all authors reviewed the manuscript draft. SMG and VPKT supervised the entire project and manuscript writing. All the authors read and approved the final version of the manuscript for submission. SMG and VPKT share equal last-author contributions. RAN and RAS share equal first-author contributions.

## Acknowledgments

The authors acknowledge all the different open-source databases and servers utilized for data mining and analyses during this study.

## Supplementary Figures

**Figure S1:**
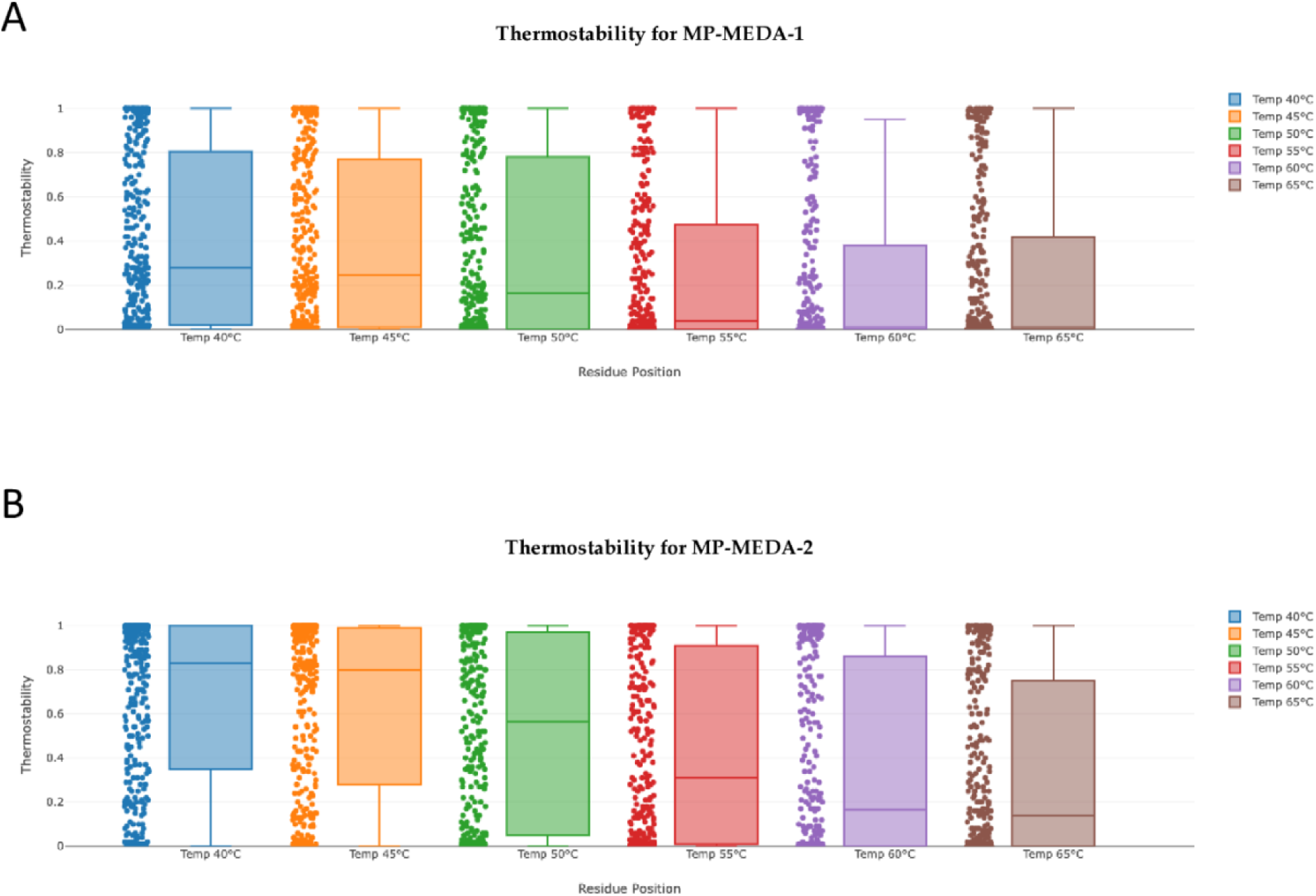
Thermostability of the designed multi-epitope diagnostic antigens. A. The thermal stability of MP-MEDA-1 at 6 different temperatures (40, 45, 50, 55, 60, and 65 °C). B. The thermal stability of MP-MEDA-2 at 6 different temperatures (40, 45, 50, 55, 60, and 65 °C).

**Figure S2:**
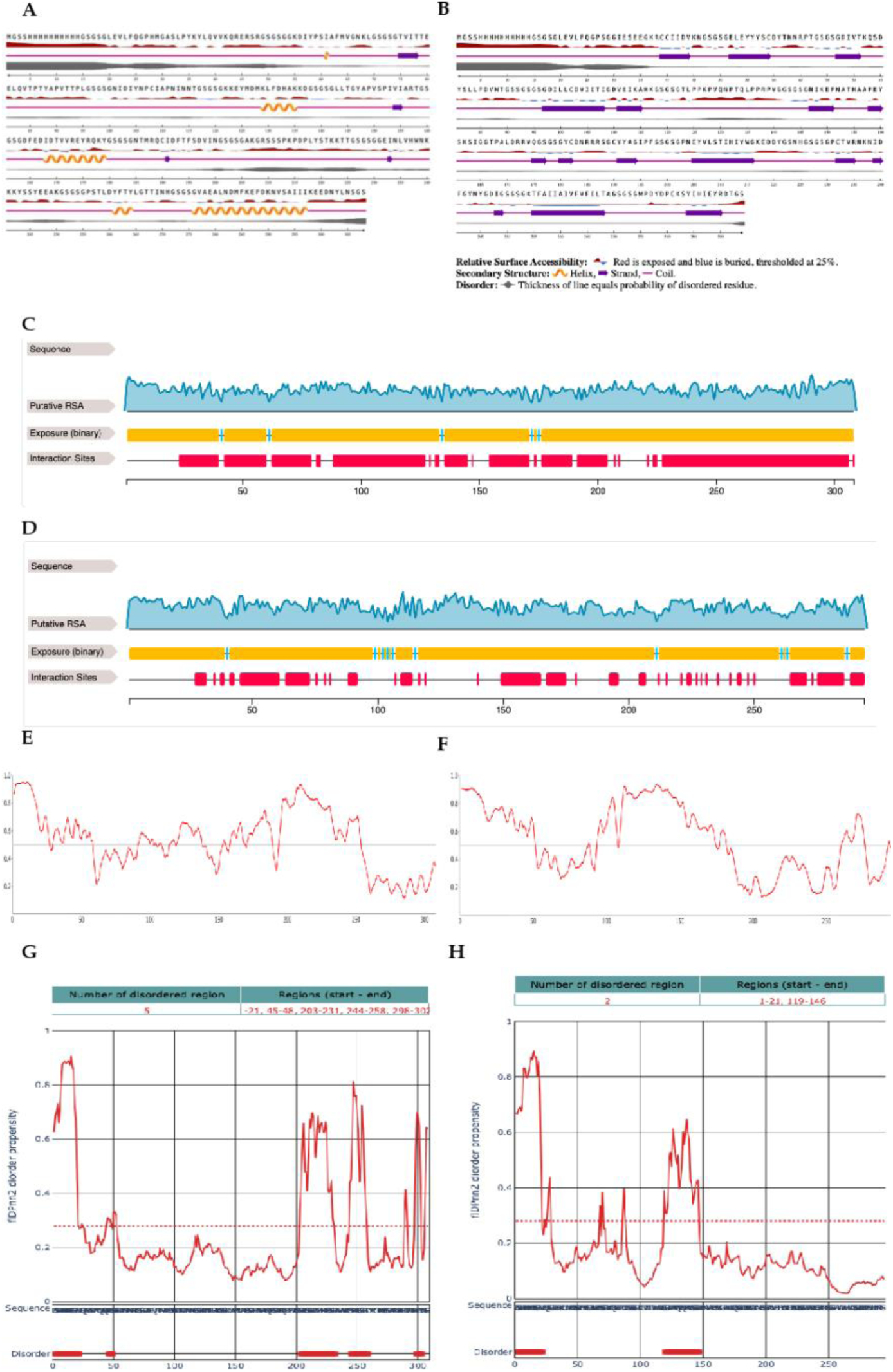
**A.** The predicted secondary structure in the MP-MEDA-1 sequence on the NetPSurf 3.0 server, showing the relative surface accessibility and intrinsic disorder. **B.** The predicted secondary structure in the MP-MEDA-2 sequence on the NetPSurf 3.0 server, showing the relative surface accessibility and intrinsic disorder. The predicted relative solvent accessibility of MP-MEDA-1 (**C**) and MP-MEDA-2 (**D**). MP-MEDA-1 is predicted to possess 303 exposed residues (yellow) and 5 buried residues (blue). MP-MEDA-2 is predicted to possess 284 exposed residues (yellow) and 10 buried residues (blue). The number of residues at the interaction sites (red regions) is 249 for MP-MEDA-1 and 144 for MP-MEDA-2. The AUIpred server intrinsic disorder plots for MP-MEDA-1 (**E**) and MP-MEDA-2 (**F),** showing the presence of disordered regions across the span of both proteins. The fIPnn2 server prediction plots for intrinsic disorder, showing the 5 disordered regions in MP-MEDA-1 (**G**) and 2 disordered regions in MP-MEDA-2 (**H**).

**Figure S3:**
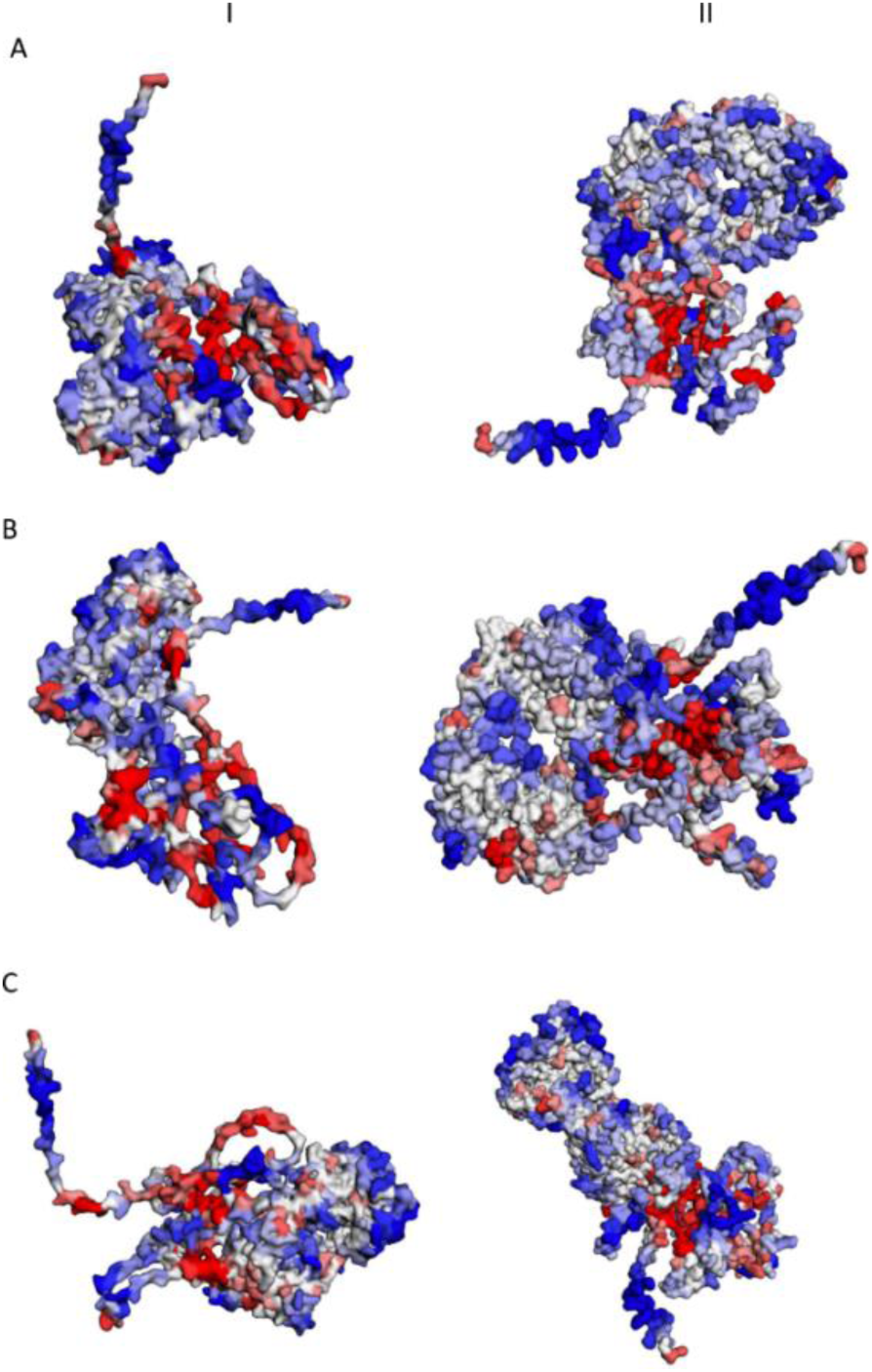
The solubility and aggregation propensity of the MEDA–Ig Fab complexes. Panels I and II show interactions of MP-MEDA-1 and MP-MEDA-2, respectively, with (A) IgA Fab, (B) IgG Fab, and (C) IgM Fab. The graphical representation model shows the soluble residues in red, the aggregation-prone residues in blue, and residues with no predicted influence shown in white.

**Figure S4:**
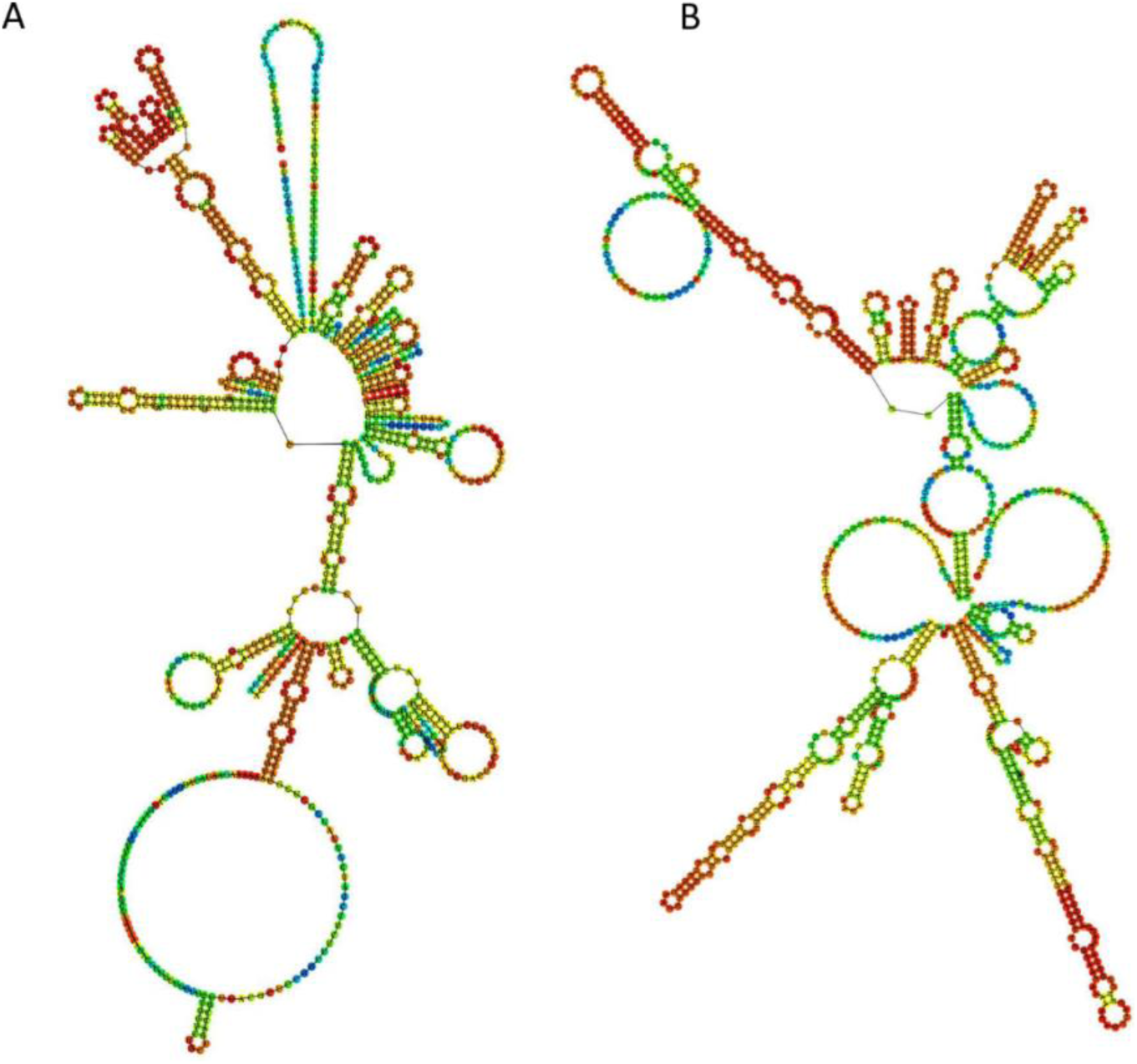
Predicted secondary structures of MP-MEDA-1 and MP-MEDA-2 mRNA sequences. The centroid secondary structures of the mRNA sequence coding for MP-MEDA-1 (A) and MP-MEDA-2 (**B**).

**Figure S5:**
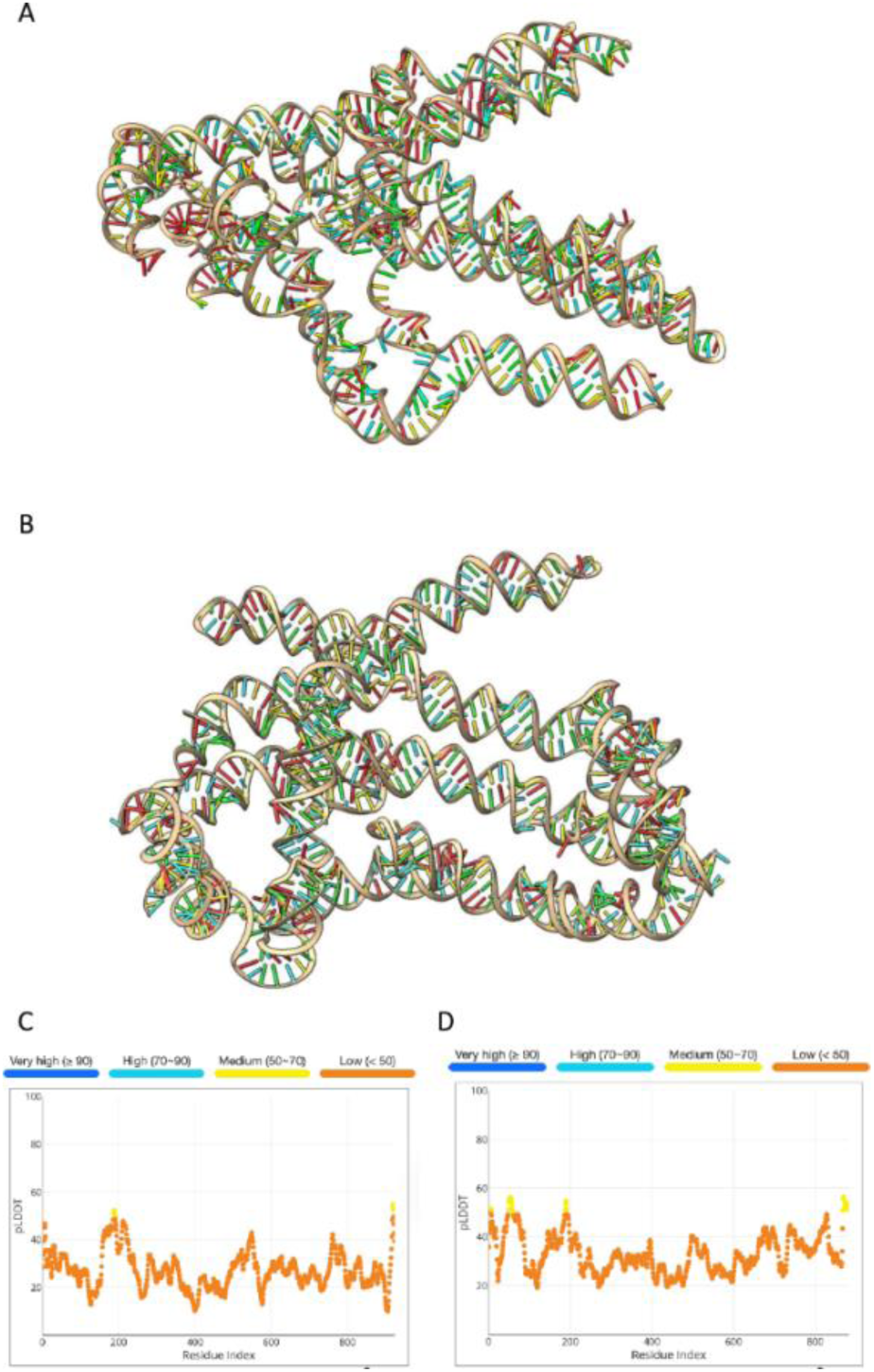
mRNA tertiary structures for MP-MEDA-1 and MP-MEDA-2. The tertiary structures of the mRNA template for MP-MEDA-1 (**A**) with a pLDDT of 27.1 and MP-MEDA-2 with a pLDDT of 33.2 (**B**) as predicted by AlphaFold3 and visualized using Chimera X (v1.9). The predicted per-residue LDDT for the MP-MEDA-1 (**C**) and MP-MEDA-2 (**D**)

**Figure S6:**
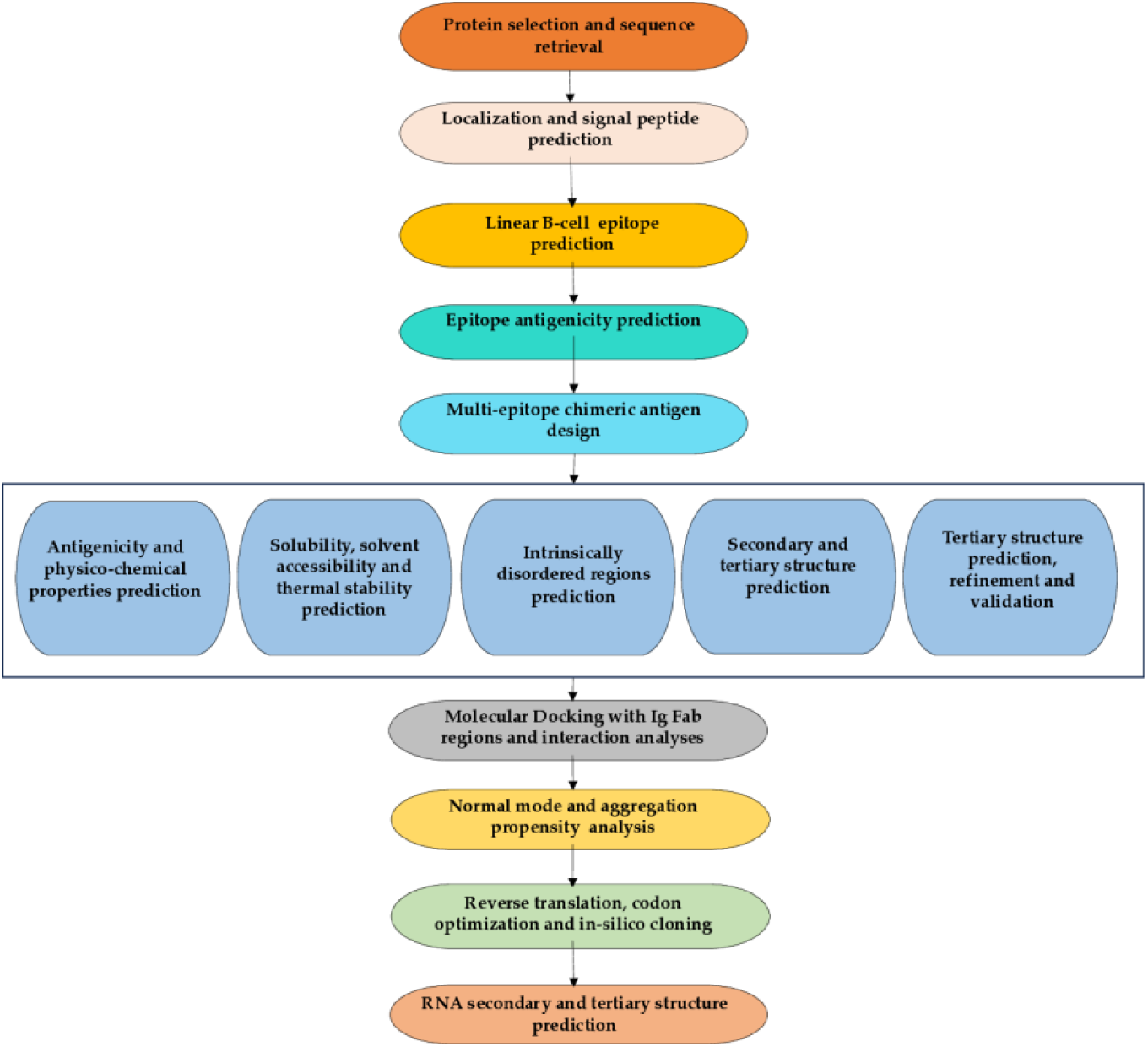
A flowchart for the study showing key steps. The entire approach used in the study comprises several phases, which began with the identification and retrieval of selected protein antigens and preliminary characterization for signal peptide and transmembrane regions. Linear B-cell epitope prediction was performed, and antigenic epitopes were selected to design two multi-epitope diagnostic antigens (MP-MEDA-1 and MP-MEDA-2). 3D modeling, refinement, and validation were performed, followed by molecular docking with the Fab regions of IgA, IgG, and IgM and normal mode analysis. Reverse translation, codon optimization, and *in silico* cloning of gene sequences coding for the designed antigens was followed by in silico transcription, secondary structure, and tertiary prediction.

## Supplementary Tables

**Table S1:** Predictable thermal stability of MP-MEDA-1 and MP-MEDA-2.

| Temperature (°C) | MP-MEDA-1 | MP-MEDA-2 |
| --- | --- | --- |
| 40 | 0.41 | 0.67 |
| 45 | 0.39 | 0.65 |
| 50 | 0.35 | 0.53 |
| 55 | 0.26 | 0.44 |
| 60 | 0.23 | 0.37 |
| 65 | 0.24 | 0.33 |

**Table S2:** Predicted discontinuous epitopes in MP-MEDA-1.

| No. | Residues | Number of residues | Score |
| --- | --- | --- | --- |
| 1 | A:M1, A:G2, A:S3, A:S4, A:H5, A:H6 | 6 | 0.968 |
| 2 | A:H8, A:H9, A:H10 | 3 | 0.968 |
| 3 | A:H14, A:G15, A:S16, A:G17, A:S18, A:G19, A:L20 | 7 | 0.926 |
| 4 | A:S210, A:S211, A:S212, A:P213, A:K214, A:P215, A:D216, A:P217, A:L218, A:Y219 | 10 | 0.922 |
| 5 | A:R44, A:E45, A:R46, A:S47, A:R48 | 5 | 0.758 |
| 6 | A:V84, A:T85, A:P86, A:T87, A:Y88, A:A89, A:P90, A:V91, A:T92, A:T93, A:P94, A:L95, A:G96, A:S97, A:G98, A:S99, A:G100, A:N101, A:I102, A:D103 | 20 | 0.738 |
| 7 | A:I297, A:K298, A:E300, A:D301, A:N302, A:Y303, A:L304, A:N305, A:S306, A:G307, A:S308 | 11 | 0.736 |
| 8 | A:N200, A:G201, A:S202, A:G203, A:S204, A:G205, A:A206, A:K207, A:G208, A:R209 | 10 | 0.732 |
| 9 | A:G49, A:S50, A:G51, A:S52, A:G53, A:G54, A:K55, A:D56 | 8 | 0.718 |
| 10 | A:S220, A:T221, A:K222, A:K223, A:T224, A:T225, A:G226, A:S227, A:G228, A:S229, A:G230 | 11 | 0.717 |
| 11 | A:A134, A:K135, A:K136, A:D137, A:G138, A:S139, A:G140, A:S141, A:G142, A:L143, A:L144, A:T145, A:G146 | 13 | 0.676 |
| 12 | A:E21, A:V22, A:L23, A:F24, A:Q25, A:G26, A:P27, A:H28 | 8 | 0.627 |
| 13 | A:N239, A:K242, A:Y243, A:S244, A:S245, A:E247, A:E248, A:A249, A:K250, A:G251, A:S252, A:G253, A:S254, A:G255,<br>A:P256, A:S257, A:T258, A:L259 | 18 | 0.617 |
| 14 | A:D167, A:D169, A:T170, A:R173, A:E174, A:R176, A:Q177, A:K178, A:Y179 | 9 | 0.598 |
| 15 | A:G180, A:S181, A:G182, A:S183 | 4 | 0.592 |
| 16 | A:G117, A:S118, A:G119, A:S120, A:G121, A:K122, A:E124 | 7 | 0.587 |

**Table S3:** Predicted discontinuous epitopes in MP-MEDA-2.

| No. | Residues | Number of residues | Score |
| --- | --- | --- | --- |
| 1 | A:M1, A:G2, A:S3, A:S4, A:H5, A:H6 | 6 | 0.98 |
| 2 | A:H7, A:H8, A:H9, A:H10, A:H11 | 5 | 0.969 |
| 3 | A:K127, A:P128, A:V129, A:Q130, A:N131, A:P132, A:T133, A:Q134, A:L135, A:P136, A:P137, A:R138, A:P139, A:V140, A:G141, A:G142, A:S143, A:G144, A:S145, A:G146, A:N147, A:I148 | 22 | 0.842 |
| 4 | A:Q78, A:S79, A:D80, A:Y81, A:S82, A:L83, A:L84, A:P85, A:D86, A:V87, A:N88, A:T89, A:G90, A:S91, A:S92, A:G93, A:S94, A:G95, A:S96, A:G97 | 20 | 0.783 |
| 5 | A:C59, A:D60, A:Y61, A:T62, A:N63, A:N64, A:R65, A:P66, A:T67, A:G68, A:S69, A:G70, A:S71, A:G72, A:D73, A:I74, A:V75, A:T76 | 18 | 0.746 |
| 6 | A:Y245, A:G246, A:D247, A:I248, A:G249, A:S250, A:G251, A:S252, A:G253, A:K254, A:T255, A:F256 | 12 | 0.696 |
| 7 | A:H12, A:H13, A:H14, A:G15, A:S16, A:G17, A:S18, A:G19 | 8 | 0.695 |
| 8 | A:V174, A:Q175, A:G176, A:S177, A:G178, A:S179, A:G180, A:Y181, A:C182, A:D183, A:N184, A:R185, A:R187, A:S188, A:G189, A:C190, A:Y191 | 17 | 0.646 |
| 9 | A:S119, A:G120, A:S121, A:G122, A:T123, A:L124, A:P125, A:P126 | 8 | 0.642 |
| 10 | A:V44, A:K45, A:N46, A:G47, A:S48, A:G49, A:S50, A:G51, A:E52, A:L53, A:E54, A:Y55 | 12 | 0.597 |
| 11 | A:L23, A:Q25, A:G26, A:P27, A:S28, A:G29, A:G30, A:I31, A:E32, A:S33, A:E34, A:K37 | 12 | 0.588 |
| 12 | A:D291, A:T292, A:G293, A:S294 | 4 | 0.553 |
| 13 | A:G228, A:S229, A:G230 | 3 | 0.536 |
| 14 | A:G270, A:S271, A:G272, A:S273, A:G274 | 5 | 0.522 |

**Table S4:** Atomic interactions in the docked complexes between MP-MEDA-1 and MP-MEDA-2 and the Fab regions of IgA, IgG, and IgM.

| Protein-protein complex | Chains | No. of interface residues | Interface area (Å<sup>2</sup>) | No. of salt bridges | No. of S-S bonds | No. of H-bonds | No. of non-bonded contacts |
| --- | --- | --- | --- | --- | --- | --- | --- |
| IgA/MP-MEDA-1 | H-L | 36:41 | 1800:1766 | 0 | 1 | 15 | 236 |
|  | Ag-H | 29:24 | 1375:1381 | 1 | 0 | 4 | 365 |
|  | Ag-L | 27:19 | 1172:1234 | 0 | 0 | 6 | 138 |
| IgG/MP-MEDA-1 | H-L | 31:38 | 1774:1782 | 2 | 10 | 14 | 189 |
|  | Ag-H | 13:14 | 638:617 | 1 | 0 | 12 | 113 |
|  | Ag-L | 14:15 | 545:492 | 1 | 0 | 11 | 81 |
| IgM/MP-MEDA-1 | H-L | 35:39 | 1906:1889 | 2 | 1 | 22 | 201 |
|  | Ag-H | 21:25 | 974:920 | 4 | 0 | 19 | 171 |
|  | Ag-L | 15:23 | 922:784 | 0 | 0 | 13 | 110 |
| IgA/MP-MEDA-2 | H-L | 37:44 | 1818:1793 | 0 | 1 | 18 | 237 |
|  | Ag-H | 21:21 | 1005:1004 | 2 | 0 | 19 | 167 |
|  | Ag-L | 11:14 | 643:589 | 0 | 0 | 9 | 55 |
| IgG/MP-MEDA-2 | H-L | 31:39 | 1796:1796 | 2 | 0 | 15 | 153 |
|  | Ag-H | 20:27 | 1098:1018 | 3 | 0 | 13 | 142 |
|  | Ag-L | 7:5 | 264:293 | 1 | 0 | 4 | 36 |
| IgM/MP-MEDA-2 | H-L | 35:38 | 1908:1905 | 2 | 1 | 21 | 203 |
|  | Ag-H | 16:15 | 658:678 | 3 | 0 | 10 | 104 |
|  | Ag-L | 7:5 | 279:311 | 1 | 0 | 7 | 37 |

Ag: Antigen, L: Light chain, H: Heavy chain

**Table S5:** Clusters predicted at the antigen-antibody complexes with the predicted interface residues.

| Protein-protein complex | Antigen-light chain interactions (clusters) | Antigen-heavy chain interactions (clusters) |
| --- | --- | --- |
| IgA/MP-MEDA-1 | 1. [['A', 'THR', 224], ['A', 'THR', 225], ['A', 'SER', 227], ['A', 'GLY', 228], ['A', 'SER', 229], ['A', 'GLY', 230], ['A', 'GLY', 231], ['A', 'GLU', 232], ['A', 'ILE', 233], ['A', 'ASN', 234], ['A', 'ASP', 260], ['A', 'TYR', 261], ['A', 'PHE', 262], ['A', 'THR', 263], ['A', 'TYR', 264], ['A', 'LEU', 265], ['A', 'GLY', 266], ['L', 'GLN', 3], ['L', 'THR', 5], ['L', 'GLN', 6], ['L', 'SER', 7], ['L', 'PRO', 8], ['L', 'SER', 9], ['L', 'SER', 10], ['L', 'PRO', 40], ['L', 'GLY', 41], ['L', 'LYS', 42], ['L', 'THR', 85], ['L', 'TYR', 87], ['L', 'PHE', 97], ['L', 'GLY', 98], ['L', 'GLY', 99], ['L', 'GLY', 100], ['L', 'THR', 101], ['L', 'ASN', 102], ['L', 'GLN', 104], ['L', 'GLU', 164]]<br>2. [['A', 'GLY', 201], ['A', 'SER', 202], ['A', 'GLY', 203], ['A', 'SER', 204], ['A', 'ALA', 206], ['A', 'LYS', 207], ['L', 'GLN', 146], ['L', 'ASN', 151], ['L', 'LEU', 153], | 1. [['A', 'HIS', 28], ['A', 'TYR', 37], ['A', 'ASP', 216], ['A', 'PRO', 217], ['A', 'LEU', 218], ['A', 'TYR', 219], ['A', 'SER', 220], ['A', 'THR', 221], ['A', 'LYS', 222], ['A', 'LYS', 223], ['A', 'THR', 225], ['A', 'GLY', 226], ['A', 'GLY', 230], ['A', 'GLY', 231], ['A', 'TYR', 264], ['A', 'LEU', 265], ['A', 'GLY', 266], ['A', 'THR', 267], ['A', 'THR', 268], ['A', 'GLY', 272], ['A', 'SER', 273], ['A', 'GLY', 274], ['A', 'SER', 275], ['A', 'GLY', 276], ['A', 'VAL', 277], ['A', 'GLU', 279], ['A', 'ALA', 280], ['A', 'ASN', 282], ['A', 'ASP', 283], ['A', 'MET', 284], ['A', 'LYS', 286], ['A', 'GLU', 287], ['A', 'LYS', 290], ['H', 'ARG', 40], ['H', 'PRO', 42], ['H', 'PRO', 43], ['H', 'GLY', 44], ['H', 'LYS', 45], ['H', 'ALA', 46], ['H', 'LEU', 47], ['H', 'GLU', 48], ['H', 'ARG', 68], ['H', 'SER', 86], ['H', 'THR', 88], ['H', 'ALA', 89], ['H', 'ALA', 90], ['H', 'THR', 92], ['H', |
|  | ['L', 'GLU', 194], ['L', 'SER', 202], ['L', 'PRO', 203],<br>['L', 'THR', 205]]<br>3. [['A', 'ASN', 270], ['A', 'HIS', 271], ['A', 'GLY', 272],<br>['L', 'ARG', 141], ['L', 'SER', 158], ['L', 'GLN', 159],<br>['L', 'GLU', 160], ['L', 'SER', 161]]<br>4. [['A', 'LYS', 214], ['A', 'ASP', 216], ['A', 'LEU', 218],<br>['L', 'SER', 113], ['L', 'ASN', 137], ['L', 'ASP', 166],<br>['L', 'LYS', 168], ['L', 'ASP', 169]]<br>5. [['A', 'LEU', 23], ['A', 'GLN', 25], ['L', 'LYS', 125], ['L',<br>'SER', 126], ['L', 'GLY', 127], ['L', 'THR', 128]] | 'VAL', 121], ['H', 'ASN', 144], ['H', 'PHE', 155],<br>['H', 'GLN', 157], ['H', 'GLU', 158], ['H', 'LEU',<br>160], ['H', 'SER', 161], ['H', 'VAL', 162], ['H',<br>'THR', 163], ['H', 'TRP', 164], ['H', 'GLU', 166],<br>['H', 'GLY', 168], ['H', 'GLY', 170], ['H', 'VAL',<br>171], ['H', 'THR', 172], ['H', 'ALA', 173], ['H',<br>'ARG', 174], ['H', 'ASN', 175], ['H', 'PRO', 177],<br>['H', 'PRO', 178], ['H', 'SER', 179], ['H', 'GLN',<br>180], ['H', 'ASP', 181], ['H', 'ALA', 182], ['H',<br>'SER', 183], ['H', 'GLY', 184], ['H', 'ASP', 185],<br>['H', 'TYR', 187], ['H', 'LEU', 193], ['H', 'THR',<br>194], ['H', 'LEU', 195]]<br>2. [['A', 'LYS', 36], ['H', 'PRO', 14], ['H', 'SER', 15]]<br>3. [['A', 'SER', 212], ['A', 'PRO', 213], ['A', 'LYS',<br>214], ['H', 'ASP', 142]]<br>4. [['A', 'ALA', 294], ['H', 'SER', 67]] |
|  |  | 5. [['A', 'ASN', 291], ['H', 'SER', 64]] |
| IgG/MP-MEDA-1 | 1. [['A', 'ILE', 199], ['A', 'ASN', 200], ['A', 'GLY', 201], ['A', 'SER', 202], ['A', 'GLY', 203], ['A', 'SER', 204], ['A', 'GLY', 205], ['A', 'LYS', 207], ['L', 'ARG', 30], ['L', 'ASN', 31], ['L', 'TYR', 32], ['L', 'LEU', 46], ['L', 'PHE', 49], ['L', 'ASP', 50], ['L', 'SER', 52], ['L', 'ASN', 53], ['L', 'GLU', 55]]<br>2. [['A', 'LYS', 68], ['A', 'LEU', 69], ['A', 'GLY', 70], ['L', 'GLN', 24], ['L', 'THR', 69], ['L', 'HIS', 70]]<br>3. [['A', 'GLY', 272], ['A', 'SER', 273], ['A', 'GLY', 274], ['A', 'SER', 275], ['L', 'ASP', 1], ['L', 'ILE', 2], ['L', 'GLN', 27], ['L', 'ASP', 28], ['L', 'GLU', 93]]<br>4. [['A', 'GLN', 25], ['A', 'HIS', 28], ['L', 'GLN', 3]] | 1. [['A', 'GLY', 203], ['A', 'SER', 204], ['A', 'GLY', 205], ['A', 'LYS', 207], ['A', 'GLY', 208], ['A', 'ARG', 209], ['A', 'SER', 210], ['A', 'SER', 211], ['A', 'SER', 212], ['A', 'PRO', 213], ['A', 'LYS', 214], ['A', 'PRO', 215], ['A', 'ASP', 216], ['A', 'LEU', 218], ['A', 'TYR', 219], ['H', 'LEU', 2], ['H', 'GLY', 26], ['H', 'GLU', 27], ['H', 'SER', 28], ['H', 'SER', 30], ['H', 'ASN', 31], ['H', 'SER', 32], ['H', 'ALA', 33], ['H', 'TYR', 34], ['H', 'ASP', 101], ['H', 'PRO', 102], ['H', 'TYR', 53], ['H', 'THR', 54], ['H', 'GLY', 55], ['H', 'THR', 73], ['H', 'ARG', 94], ['H', 'ILE', 97], ['H', 'VAL', 98], ['H', 'TRP', 100]] |
| IgM/MP-MEDA-1 | 1. [['A', 'GLY', 146], ['A', 'TYR', 147], ['A', 'PRO', 149], ['A', 'ASN', 185], ['A', 'ARG', 188], ['A', 'GLN', 189], ['A', 'CYS', 190], ['A', 'ASP', 192], ['A', 'HIS', 237], ['A', 'TRP', 238], ['A', 'ASN', 239], ['A', 'LYS', 240], ['A', 'LYS', 241], ['A', 'LYS', 242], ['A', 'GLY', 255], | 1. [['A', 'ASN', 239], ['A', 'LYS', 242], ['A', 'TYR', 243], ['A', 'SER', 252], ['A', 'SER', 254], ['A', 'GLY', 255], ['A', 'SER', 257], ['A', 'THR', 258], ['H', 'LEU', 550], ['H', 'TYR', 559], ['H', 'PHE', 601], ['H', 'TYR', 602], ['H', 'ASP', 603], ['H', |
|  | ['A', 'PRO', 256], ['A', 'SER', 257], ['A', 'THR', 258],<br>['A', 'LEU', 259], ['A', 'ASP', 260], ['A', 'PHE', 262],<br>['L', 'ASP', 1], ['L', 'ILE', 2], ['L', 'GLN', 3], ['L', 'THR',<br>5], ['L', 'ARG', 24], ['L', 'SER', 26], ['L', 'GLN', 27],<br>['L', 'SER', 28], ['L', 'SER', 30], ['L', 'SER', 31], ['L',<br>'TYR', 32], ['L', 'TYR', 49], ['L', 'ALA', 50], ['L',<br>'ALA', 51], ['L', 'SER', 52], ['L', 'SER', 65], ['L', 'GLY',<br>66], ['L', 'SER', 67], ['L', 'SER', 91], ['L', 'TYR', 92],<br>['L', 'SER', 93], ['L', 'ALA', 94], ['L', 'PRO', 95]]<br>2. [['A', 'SER', 52], ['A', 'GLY', 53], ['A', 'GLY', 54], ['A',<br>'LYS', 55], ['A', 'ILE', 57], ['L', 'PRO', 8], ['L', 'SER',<br>10], ['L', 'LEU', 11], ['L', 'SER', 12], ['L', 'THR', 20],<br>['L', 'THR', 22], ['L', 'GLU', 105], ['L', 'LYS', 107], ['L',<br>'ARG', 108], ['L', 'THR', 109], ['L', 'VAL', 110], ['L',<br>'TYR', 140], ['L', 'ARG', 142], ['L', 'GLU', 143]] | 'PRO', 604], ['H', 'THR', 605], ['H', 'ALA', 606],<br>['H', 'PRO', 607]] |
| IgA/MP-MEDA-2 | 1. [['A', 'SER', 177], ['A', 'GLY', 178], ['A', 'SER', 179],<br>['A', 'GLY', 180], ['A', 'TYR', 181], ['A', 'TYR', 243],<br>['A', 'ASN', 244], ['A', 'TYR', 245], ['A', 'GLY', 246],<br>['A', 'ASP', 247], ['A', 'ASP', 291], ['A', 'GLY', 293],<br>['A', 'SER', 294], ['L', 'SER', 126], ['L', 'GLY', 127], | 1. [['A', 'SER', 143], ['A', 'GLY', 144], ['A', 'SER',<br>145], ['A', 'GLY', 146], ['A', 'ASN', 147], ['A',<br>'ILE', 148], ['A', 'LYS', 149], ['A', 'GLU', 150],<br>['A', 'ASN', 244], ['A', 'TYR', 245], ['A', 'GLY',<br>246], ['A', 'ASP', 247], ['A', 'ILE', 248], ['A',<br>'GLY', 249], ['A', 'SER', 250], ['A', 'GLY', 251],<br>['A', 'SER', 252], ['A', 'GLY', 253], ['A', 'LYS',<br>254], ['A', 'PHE', 256], ['A', 'TYR', 289], ['A', |
|  | ['L', 'THR', 128], ['L', 'ALA', 152], ['L', 'LEU', 153],<br>['L', 'GLN', 154], ['L', 'SER', 155], ['L', 'GLY', 156],<br>['L', 'ASN', 157], ['L', 'SER', 158], ['L', 'GLN', 159],<br>['L', 'GLU', 160], ['L', 'THR', 179], ['L', 'LEU', 180],<br>['L', 'SER', 181], ['L', 'ASP', 184], ['L', 'LYS', 187],<br>['L', 'HIS', 188]]<br>2. [['A', 'GLY', 141], ['A', 'GLY', 142], ['A', 'SER', 143],<br>['L', 'ARG', 141], ['L', 'VAL', 162], ['L', 'GLU', 164]] | 'ASP', 291], ['A', 'THR', 292], ['A', 'GLY', 293],<br>['A', 'SER', 294], ['H', 'PRO', 43], ['H', 'GLY', 44],<br>['H', 'ALA', 89], ['H', 'THR', 92], ['H', 'LEU', 118],<br>['H', 'THR', 120], ['H', 'VAL', 121], ['H', 'SER',<br>122], ['H', 'ALA', 124], ['H', 'SER', 125], ['H',<br>'PRO', 126], ['H', 'THR', 127], ['H', 'SER', 128],<br>['H', 'LYS', 130], ['H', 'GLN', 152], ['H', 'GLY',<br>153], ['H', 'PHE', 155], ['H', 'GLN', 157], ['H',<br>'GLU', 158], ['H', 'PRO', 177], ['H', 'PRO', 178],<br>['H', 'SER', 179], ['H', 'GLN', 180], ['H', 'ASP',<br>181], ['H', 'ALA', 182], ['H', 'SER', 183], ['H',<br>'GLY', 184], ['H', 'ASP', 185], ['H', 'LEU', 186],<br>['H', 'TYR', 187], ['H', 'THR', 188]] |
| IgG/MEDA-2 | 1. [['A', 'SER', 28], ['A', 'GLY', 29], ['A', 'GLY', 30], ['A',<br>'ILE', 31], ['A', 'GLU', 32], ['A', 'SER', 50], ['A', 'GLY',<br>51], ['A', 'GLU', 52], ['A', 'LEU', 53], ['A', 'TYR', 56],<br>['L', 'ASP', 122], ['L', 'GLU', 123], ['L', 'GLN', 124],<br>['L', 'LYS', 126], ['L', 'SER', 127], ['L', 'THR', 129],<br>['L', 'LYS', 183], ['L', 'GLU', 187]] | 1 [['A', 'ASP', 80], ['A', 'TYR', 81], ['A', 'SER', 82],<br>['A', 'LEU', 83], ['A', 'LEU', 84], ['A', 'PRO', 85],<br>['H', 'GLN', 3], ['H', 'GLN', 5], ['H', 'GLU', 6], ['H',<br>'SER', 7], ['H', 'GLY', 8], ['H', 'SER', 23], ['H',<br>'SER', 25], ['H', 'SER', 74], ['H', 'LYS', 75], ['H',<br>'ASN', 76], ['H', 'GLN', 77]]<br>2 [['A', 'ASN', 46], ['A', 'SER', 50], ['A', 'GLU', 52],<br>['A', 'LEU', 53], ['A', 'GLU', 54], ['A', 'TYR', 55],<br>['A', 'TYR', 56], ['A', 'TYR', 57], ['A', 'SER', 58],<br>['A', 'ILE', 105], ['A', 'THR', 107], ['A', 'GLY',<br>109], ['A', 'ASP', 110], ['A', 'GLU', 112], ['A',<br>'LYS', 114], ['H', 'LEU', 11], ['H', 'LYS', 13], ['H',<br>'ALA', 114], ['H', 'SER', 115], ['H', 'THR', 116],<br>['H', 'LYS', 117], ['H', 'GLY', 118], ['H', 'PRO',<br>119], ['H', 'SER', 120], ['H', 'VAL', 121], ['H',<br>'PHE', 122], ['H', 'LYS', 143], ['H', 'ASP', 144],<br>['H', 'SER', 203], ['H', 'ASN', 204], ['H', 'THR', |
|  |  | 205], ['H', 'LYS', 206], ['H', 'VAL', 207], ['H', 'ASP', 208], ['H', 'LYS', 209]]<br>3 [['A', 'ASP', 60], ['A', 'THR', 62], ['H', 'LYS', 210]]<br>4 [['A', 'LYS', 77], ['H', 'SER', 156], ['H', 'GLY', 157], ['H', 'ALA', 158]] |
| IgM/MEDA-2 | 1. [['A', 'LYS', 77], ['L', 'ASP', 1], ['L', 'ALA', 94], ['L', 'PRO', 95]]<br>2. [['A', 'GLU', 52], ['A', 'LEU', 53], ['A', 'TYR', 56], ['L', 'TYR', 49], ['L', 'GLN', 55], ['L', 'SER', 56]]<br>[['A', 'LYS', 37], ['A', 'SER', 58], ['A', 'CYS', 59], ['A', 'ASP', 60], ['A', 'THR', 62], ['L', 'SER', 28], ['L', 'SER', 30], ['L', 'TYR', 32]] | 1. [['A', 'GLU', 52], ['A', 'LEU', 53], ['A', 'GLU', 54], ['A', 'TYR', 55], ['A', 'TYR', 56], ['A', 'TYR', 57], ['A', 'SER', 58], ['A', 'SER', 79], ['A', 'ASP', 80], ['A', 'TYR', 81], ['A', 'ILE', 105], ['A', 'THR', 107], ['A', 'GLY', 109], ['A', 'ASP', 110], ['A', 'GLU', 112], ['A', 'LYS', 114], ['H', 'THR', 528], ['H', 'SER', 530], ['H', 'GLY', 531], ['H', 'TYR', 532], ['H', 'TYR', 553], ['H', 'ASP', 554], ['H', 'GLU', 555], ['H', 'SER', 556], ['H', 'ASN', 557], ['H', 'LYS', 558], ['H', 'TYR', 559], ['H', 'LYS', 600], ['H', 'PHE', 601], ['H', 'TYR', 602], ['H', 'ASP', 603]]<br>2. [['A', 'LYS', 77], ['H', 'ASP', 562], ['H', 'LYS', 565]] |

